# Experience-dependent structural plasticity in the adult brain: How the learning brain grows

**DOI:** 10.1101/2020.09.01.277467

**Authors:** Silvio Schmidt, Sidra Gull, Karl-Heinz Herrmann, Marcus Boehme, Andrey Irintchev, Anja Urbach, Jürgen R. Reichenbach, Carsten M. Klingner, Christian Gaser, Otto W. Witte

**Author notes:** These authors contributed equally (Silvio Schmidt, Sidra Gull). **Correspondence to:** Otto W. Witte, Hans Berger Department of Neurology, Jena University Hospital, Am Klinikum 1, D07747 Jena, Germany.

## Abstract

Volumetric magnetic resonance imaging studies have shown that intense learning can be associated with grey matter volume increases in the adult brain. The underlying mechanisms are poorly understood. Here we used monocular deprivation in rats to analyze the mechanisms underlying use-dependent grey matter increases. Optometry for quantification of visual acuity was combined with volumetric magnetic resonance imaging and microscopic techniques in longitudinal and cross-sectional studies. We found an increased spatial vision of the open eye which was associated with a transient increase in the volumes of the contralateral visual and lateral entorhinal cortex. In these brain areas dendrites of neurons elongated, and there was a strong increase in the number of spines, the targets of synapses, which was followed by spine maturation and partial pruning. Astrocytes displayed a transient pronounced swelling and underwent a reorganization of their processes. The use-dependent increase in grey matter corresponded predominantly to the swelling of the astrocytes. Experience-dependent increase in brain grey matter volume indicates a gain of structure plasticity with both synaptic and astrocyte remodeling.

**Highlights:**

- Perception learning causes a transient increase in brain grey matter volume detectable by MRI.
- This learning results in pronounced changes of neuronal dendrites and an increase in the number of dendritic spines.
- Structural neuronal plasticity is associated with a reorganization and transient swelling of astrocytes.
- Brain volume and astrocyte volume return to baseline post-learning, with a persistent increase in the number of mature spines.

## 1. Introduction

The brain preserves the ability of experience-dependent plasticity throughout life. One aspect of this phenomenon is synaptic plasticity, i. e. the ability to modulate synaptic function and form new and/or eliminate existing synapses (Holtmaat and Svoboda, 2009). The adaptive brain responses possibly also include other structural alterations as suggested by relatively large-scale increases in regional estimates of brain volume detectable by magnetic resonance imaging (MRI) (Lindenberger et al., 2017). Local and transient “temporal” macroscopic changes have been linked with learning in visuomotor tasks; they renormalized as volunteers reached expert performance (Draganski et al., 2004; Driemeyer et al., 2008; Taubert et al., 2010). Although not analyzed longitudinally, short-term macroscopic changes have also been observed after extensive training of golf (Bezzola et al., 2011), preparation for an academic exam (Draganski et al., 2006), learning a foreign language (Legault et al., 2018), post-stroke rehabilitation (Sampaio-Baptista et al., 2018) and also in the course of improved tactile discrimination after a single intervention by repetitive somatosensory stimulation (Schmidt-Wilcke et al., 2018). Persisting alterations in brain structure have been documented in relation to professional challenges. London cab drivers, processing well-trained spatial memory, have posterior hippocampi that are larger than those of the general population (Maguire et al., 2000). Professional musicians with highly complex motor and auditory skills show increased grey matter volume in several brain areas as compared with amateur musicians and non-musicians (Gaser and Schlaug, 2003).

The cellular basis of structural brain plasticity detectable at the MRI level is not well understood (Wenger et al., 2017). Changes in intracortical myelin (Keiner et al., 2017; Kougioumtzidou et al., 2017; McKenzie et al., 2014), enhanced neurogenesis (van Praag et al., 1999), increased spine density in neuronal dendrites (Keifer et al., 2015) and alterations in astrocyte volume (Kleim et al., 2007; Woo et al., 2018) have been suggested to contribute to plasticity in different experimental settings (Zatorre et al., 2012). In the present study, we used monocular deprivation (MD) in rats, a well-established paradigm in brain plasticity research (Prusky et al., 2006; Sato and Stryker, 2008), to analyse experience-dependent changes in macroscopic brain structure in a longitudinal fashion by using MRI and, for the first time, combined the longitudinal experiments with cross-sectional cell-morphological investigations on the microscopic level. MD of one eye gradually enhances visual acuity and contrast sensitivity of the open eye, which indicates visual perception learning with involvement of the visual cortex as shown by Prusky et al. (2006).

Using optometry and serial MRI, we found that enhanced visual experience in the pathway supplied by the open eye is linked with a transient volume increase in the contralateral primary binocular visual cortex (V1B) and also in the contralateral lateral entorhinal cortex (LEnt). Neurons in these brain areas display dendrite and spine plasticity; newly generated spines partially mature and persist beyond the learning period. The neuronal plasticity is associated with a transient massive astrocytic rearrangement, which is the main cause for the transient brain volume increase recorded by MRI. Neither neurogenesis nor gliogenesis contribute to this type of plasticity in the adult cortex. The deprived V1B contralateral to the closed eye, which has been shown to undergo compensatory homeostatic plasticity (Hofer et al., 2006), does not display a change in grey matter or in astrocyte volume.

## 2. Methods and Materials

### 2.1 Experimental design

All experiments were approved by the Governmental Animal Care and Use Committee (Thüringer Landesamt für Lebensmittelsicherheit und Verbraucherschutz; registry number: 02-048/08). Experiments were performed on 2 months old male albino Wistar rats (RccHan:WIST), housed in standard cages (5 animals per cage) on a 12 h light/dark cycle with food and water ad libitum. Interventions on living animals were performed during the light cycle.

The study consists of longitudinal (Fig. 1A) and cross-sectional experiments (Fig. 3A) to analyse brain plasticity induced by MD (a change in visual experience with loss of stereoscopic vision by closure of one eye-lid). In longitudinal experiments (Fig. 1A), visual function as well as macroscopic brain morphology were repetitively assessed *in vivo* over time by using optometry and MRI. 2 days before MD, animals were subjected to optometry to determine the baseline visual function. MD was induced on day 0 3-4 hours after obtaining an MRI. Control animals received the same procedures without MD. For the assessment of macroscopic brain morphology, datasets were collected and analyzed from n = 23 controls and n = 24 MD-animals.

**Figure 1.**
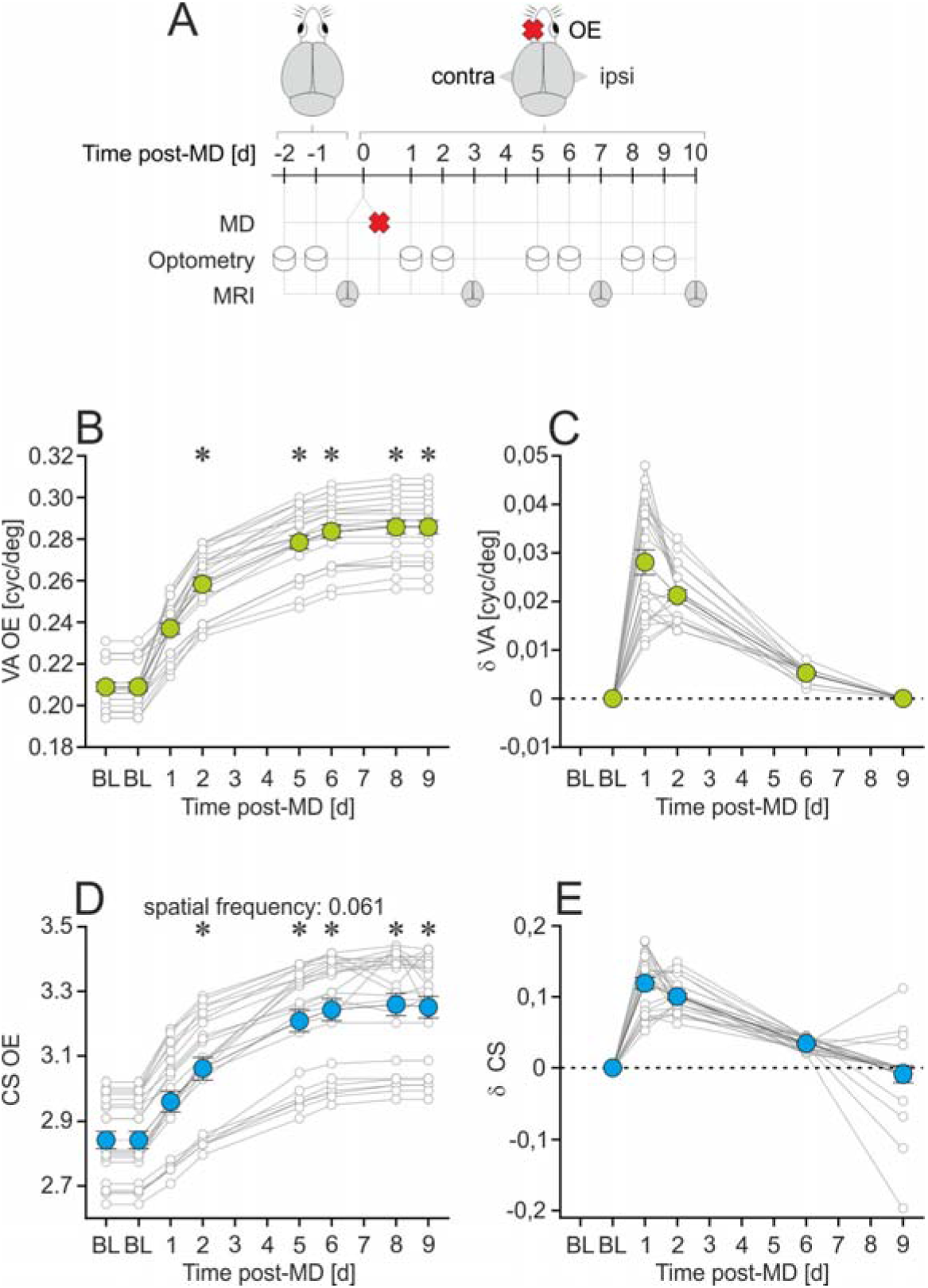
MD-induced visual gain. **(A)** Longitudinal study design: MD-induced changes in open eye (OE) visual function as well as brain macro-structure were analyzed by optometry and MRI. **(B, D)** Visual acuity (VA) and contrast sensitivity (CS, at spatial frequency of 0.061 cyc/deg) increased and reached saturation at day 8-9 post-MD (n=24). **(D, F)** Daily visual improvement rates (δVA, δCS) were highest immediately post-MD. Data are presented as the mean ± SEM. * p < 0.001 vs. BL, RM ANOVA on Ranks. For detailed values and statistics, see supplementary table S1.

In cross-sectional experiments, MD-animals received the same procedures as in the longitudinal ones, except for MRI-scans. At day 3 or 10 after MD, animals were sacrificed for microscopic analyses. For each condition, datasets were collected and analyzed from n = 5-10 animals.

### 2.2 MD

The left eye was deprived (closed) under isoflurane anesthesia (2.5 %, O2:N2O=20:40 l/h) by suturing the eyelid (Greifzu et al., 2011). Stitches were checked daily and animals which reopened the eye during the experiment were excluded from further analyses.

### 2.3 Optometry

Visual acuity (VA) and contrast sensitivity (CS) was examined by optometry as described by Prusky et al. (2004). In brief, freely moving animals were exposed to moving vertical sine-wave grating stimuli of various spatial frequencies and contrasts which they track as long as they sense it by distinct head movements (optokinetic response). For assessment of VA, spatial frequency (cycles per degree, [cyc/deg]) at 100 % contrast was increased by the experimenter until the frequency-threshold of tracking behavior was determined. CS was assessed by decreasing the contrast of the grating at five different spatial frequencies (0.044, 0.061, 0.089, 0.119, 0.15 cyc/deg) until the contrast-threshold of tracking behavior was determined. CS at a spatial frequency was calculated as a Michelson contrast based on the screen’s luminance (maximum-minimum) / (maximum+minimum). Because only motion of the grating in the temporal-to-nasal direction provokes tracking, MD of the left eye abolishes the tracking in the clockwise direction. In control animals without MD, values of VA and CS were examined separately for each eye and averaged per animal. Daily improvement rates of VA and CS (δVA, δCS) were calculated between days where optometry was performed immediately consecutive without gaps.

### 2.4 MRI

Brain MRI was performed using a clinical 3T whole body scanner (Magnetom TIM Trio, Siemens Medical Solutions, Erlangen, Germany) equipped with a dedicated rat head volume resonator using a linearly polarized Litz design (Doty Scientific Inc., Columbia, SC, USA) (Haenold et al., 2012; Herrmann et al., 2012). Freely breathing animals were anesthetized by isoflurane (1.7 % in oxygen, 1.5 l/min). T2-weighted images were obtained by using a 3D SPACE sequence (Sampling Perfection with Application-Optimized Contrasts Using Different Flip Angle Evolutions, Siemens Healthcare, Erlangen Germany) with isotropic resolution of 0.33 mm^3^ (matrix 192 × 130 × 96, FOV 64 × 43 × 32 mm, bandwidth: 145 Hz/px, TE: 352 ms, TR: 2500 ms, flip angle mode: ‘T2var’, echo spacing: 10.7 ms, turbo factor: 67, Partial Fourier: 7/8 in both phase encode directions, two repetitions averaged to a single magnitude image with TA = 14 min). From each animal, four brain images were acquired at baseline (BL, MRI-1) as well as repetitively on days 3, 7 and 10 thereafter (MRI 2-4).

### 2.5 Deformation based morphometry (DBM)

Temporal changes of brain morphology were analyzed by using the DBM tool established and described in detail by Gaser et al. (2012). The DBM tool is implemented in the MATLAB software package SPM8 and uses two nonlinear registration techniques that differ in terms of dimensionality. It works in the space of a customized template brain which was landmark-based transformed into the coordinates of the Paxinos Atlas (Paxinos and Watson, 2005). This allows for the anatomical assignment of each voxel to brain regions defined in the widely used atlas. In brief, repetitively acquired MR-images (MRI 2-4) of individual datasets (n = 24 from the MD-group, n = 23 from controls) were rigidly registered to their own BL-image (MRI 1) for positional correction and subsequently, local deformations were introduced by highdimensional nonlinear normalization to minimize local morphological differences between the image pairs. Thereafter, morphological differences were encoded in 3D deformation fields consisting of specific displacement vectors in each voxel (Jacobian determinants) which describe the volume changes between the image pairs (local shrinkage or swelling compared to BL). Finally, all intra-pair deformations were nonlinearly transformed to our customized template brain by using the same normalization parameters needed to do the same with the BL-image. For this lower-dimensional normalization, we used the default spatial normalization implemented in MATLAB software package SPM8. The displacement vectors were smoothed with a Gaussian kernel with FWHM of 0.8 mm and used for further statistical analysis.

To analyze qualitative interactions between the groups, we applied a general linear model (GLM) with an RM ANOVA (threshold of p < 0.05, corrected for family-wise error [FWE] or at an uncorrected threshold of p < 0.001) as design and tested our hypotheses using a t-statistics within this GLM. This strategy is implemented in SPM8 as flexible factorial model and we used the factors “subject”, “group” and “time” and tested for an interaction between group and time. First, we tested for brain regions where volume changes in the MD-group decreased over time in contrast to controls (MD: MRI 2 > MRI 4 vs. Control: MRI 2 < MRI 4). This comparison was used to detect where the MD induces an initial swelling, a late shrinkage or a mixture of both.

Furthermore, we tested the same approach vice versa (MD: MRI 2 < MRI 4 vs. Control: MRI 2 > MRI 4) to detect MD-induced late swellings or initial shrinkages. For quantification, the MD-induced percental volume changes were averaged per cluster and time-point (cluster threshold: p < 0.001, uncorrected, cluster cut-off: 40 voxel) and normalized to controls.

Furthermore, volume changes were analyzed a priori in V1B and LEnt as defined according to the Paxinos Atlas (Gaser et al., 2012) to check whether the hypothesis free approach could be verified by an independent analysis. The Jacobian determinants were averaged per area and the differences between the groups were compared for the time-point 3 days after MD by using the student-t test.

### 2.6 50⍰-Bromodeoxyuridine (BrdU) labeling

To label proliferating cells, the thymidine analogous BrdU, (Sigma-Aldrich, St Louis, MO, USA) was injected intra-peritoneally (100⍰mg/kg body weight; dissolved in 0.9% saline) every 12 hours for 3 consecutive days (Urbach et al., 2015). The first injection was performed immediately after MD. The animals were sacrificed 12 hours after the last injection.

### 2.7 Immunohistochemistry

Animals were transcardially perfused under isoflurane anaesthesia with 0.1 M phosphate-buffered saline (4°C, pH 7.4) followed by the same solution containing 4 % formaldehyde. Brains were removed and post-fixed overnight in the same fixative, cryoprotected in 0.1 M phosphate-buffered saline containing 10 % sucrose for 24 h and 30 % sucrose until they sank (each at 4°C), snap frozen in methyl butane at −30 °C, stored at −80 °C and sectioned into 40 μm thick coronal slices. For 3, 3’-diaminobenzidine immunostaining, endogenous peroxidases were quenched with hydrogen peroxide and non-specific binding was blocked with normal serum from the species in which the secondary antibody was raised. Primary antibodies against the activity-regulated cytoskeleton-associated protein (ARC; guinea pig, Synaptic system), the glial fibrillary acidic protein (GFAP; mouse, Chemicon), the S100 calcium-binding protein B (S100B; rabbit, Synaptic system), and BrdU (mice, AbD Serotec) were detected with biotinylated secondary antibodies (Jackson) and the Vector-Elite ABC kit (Vector Laboratories, Burlingame, CA). Finally, sections were mounted on glass slides, air dried, covered with Entellan (Merck, Darmstadt, Germany) and coded before blinded quantitative analysis.

### 2.8 Golgi staining

Neuronal staining was done by using the FD Rapid GolgiStain™ Kit (FD Neurotechnolgies, USA) according to manufacturer’s instructions. Briefly, animals were decapitated under isoflurane anaesthesia, brains were submerged in a 1:1 mixture of solution A and B (3 weeks, at room temperature) followed by immersion in solution C for 2 days at 4°C. Then, brains were snap frozen in methylbutane at −30°C, stored at −80°C, sectioned into 150 μm thick coronal slices on a cryostat (Leica CM3050 S, Germany), mounted on Superfrost slides (Thermo Scientific, USA), stained in solutions D and E for 10 minutes, covered with Entellan (Merk, Darmstadt, Germany) and coded before quantitative analysis.

For astrocytic staining, the manufacturer’s protocol was modified as described by Gull et al. (2015). Briefly, isoflurane anesthetized animals were transcardially perfused with 4% formaldehyde in 0.1 M phosphate buffer (pH 7.4), brains were post-fixed for 4 days in the same fixative additionally containing 8% glutaraldehyde, submerged in a 1:1 mixture of solution A and B for 3 days at 26°C, transferred into solution C for 7 days at 4°C, sectioned on a cryostat (Leica CM3050 S, Germany) into 150μm coronal slices, mounted on Superfrost slides (Thermo Scientific, USA), stained in solutions D and E for 10 minutes, covered with Entellan (Merk, Darmstadt, Germany) and coded before quantitative analysis.

### 2.9 Determination of regions of interest (ROI)

For microscopic analyses on brain slices, ROIs (V1B, LEnt) were defined in the space of the Paxinos atlas (Paxinos and Watson, 2005) according to the location of significant DBM-clusters, each in the hemisphere ipsilateral to the open eye (ipsi-OE) and contralateral to the opne eye (contra-OE). The auditory cortex (AU) was included as a control region in all microscopic analyses. Slices were selected from Bregma −5.0 to −6.0 to analyse the LEnt and the AU and from Bregma −6.5 to −7.5 to analyse V1B. Cortical layers were defined according to Zilles (1985) and analyses were performed in V1B in layer III/IV, in LEnt in layer II/III and AU in layer II/III. All microscopic analyses were performed blindly with respect to the experimental group (MD, control, time post-MD/sham).

### 2.10 Stereological estimation of cell density

Cell density was estimated by applying the optical fractionator method (Raslan et al., 2014) using the Stereo Investigator software 8.1 (MicroBrightField Europe, Magdeburg, Germany) and a light microscope (Axioscope 2 mot plus, Zeiss, Oberkochen, Germany) equipped with a Plan Neofluar 40x objective (Zeiss), a motorized stage (Zeiss) and a digital camera (CX 9000, MicroBrightField). Dissectors for cell-counting were placed randomly in two slices per ROI. BrdU+ cells were counted in n = 6 control and n = 7 MD-3d animals. S100B+ cells were counted in n = 8 control, n = 7 MD-3d and n = 8 MD-10d animals. ARC+ cells were counted in n = 10 control, n = 6 MD-3d and n = 6 MD-10d animals.

### 2.11 Quantification of pyramidal dendritic length and spine frequency

Analyses were performed on Golgi stained cells using the NeuroLucida 8 software (MicroBrightField Europe, Williston, VT, USA) and a light microscope (Axioscope 2 mot plus, Zeiss, Oberkochen, Germany) equipped with a Plan Neofluar 100x objective (Zeiss), a motorized stage (Zeiss) and a digital camera (CX 9000, MicroBrightField). The length of basal dendrites from cells with pyramidal shape was measured on-line by 3-dimensional tracing of n = 2-3 dendrites per neuron from n = 2-6 neurons per ROI in n = 7 control, n = 6 MD-3d and n = 7 MD-10d animals. Only dendrites (separately for 1^st^ order segments and 2^nd^-3^rd^ order segments) in which the origin and the end of the segments was clearly identifiable by branching points were analyzed. As the Golgi method randomly stains only a limited number of cells, in some ROIs we did not find at least two pyramidal cells fulfilling the analysis criteria. Such ROIs were excluded from analysis. In total, n = 945 dendritic segments from n=339 neurons were analyzed. In the same animals, but using a different pool of cells, the spine frequency (equivalent to line number density) was quantified on 2^nd^-3^rd^ order basal pyramidal dendritic segments which were completely traceable from their origin to their end following the criteria introduced by Kassem et al. (2013): they were classified as small (≤ 1μm), medium (1-1.5μm), larger (1.5-4.5μm) spines on the basis of length, and as mushroom or spiny on the basis of shape. 2-3 dendritic segments per neuron from 2-6 neurons per ROI were analysed. This summed up to a total of 24,743 spines from n = 336 pyramidal neuron. Spine frequency was normalized to 100μm of dendrite.

### 2.12 Quantification of astrocyte morphology

The complexity of the astrocytic cytoskeleton was analyzed from GFAP-stained cells of stellate shape using the NeuroLucida 8 software (MicroBrightField Europe, Williston, VT, USA) and a light microscope (Axioscope 2 mot plus, Zeiss, Oberkochen, Germany) equipped with a Plan Neofluar 100x objective (Zeiss), a motorized stage (Zeiss) and a digital camera (CX 9000, MicroBrightField). First, the mesh of GFAP-stained processes was 3 dimensionally constructed from seven cells per ROI (randomly selected from 2-4 slices) and then analyzed by the Sholl’s concentric circle method (Dall’Oglio et al., 2008; Sholl, 1953) in n = 6 control, n = 6 MD-3d and n = 6 MD-10d animals. Primary processes were quantified by counting these ones extending directly from the soma of the same cell.

The territorial volume was analyzed on Golgi stained cells of spherical shape following the procedure described by Grosche et al. (2013) by using a light microscope (Axioscope 2 mot plus, Zeiss, Oberkochen, Germany) equipped with a Plan Neofluar 100x objective (Zeiss), a motorized stage (Zeiss) and a digital camera (CX 9000, MicroBrightField). The Golgi technique delineates every fine detail of the cells and the cell territorial volume was defined as the space over which all elaborations extend. Based on optical slices (gap: 2 μm), the area of maximal cell extension was determined from 30 cells per ROI (randomly selected from 2-4 slices) by using Image J, treated as circular and used to calculate the volume of theoretical “astrocytic globes” in n = 5 control, n = 7 MD-3d and n = 7 MD-10d animals. The soma volume of astrocytes was analyzed in the same manner on S100B stained cells in ROIs from n = 10 control, n = 6 MD-3d and n = 6 MD-10d animals.

### 2.13 Statistical analysis

All animals or brain samples that fulfilled proper experimental criteria during the experimental procedures were included in the analysis. For microscopic analyses the investigators were blinded with respect to group to which the samples belonged (control vs. intervention). Statistics was performed by using Sigma plot or SPSS and a student-t test, repeated measures analysis of variance (RM ANOVA) on ranks with Tukey’s post-hoc test, one-way ANOVA with Tukey’s post-hoc test or Mann-Whitney rank sum test. Statistics for the quantification of pyramidal dendritic length and spine frequency and for the complexity of the astrocytic cytoskeleton is based on the number of cells per group. Statistics for the quantification of astrocytic soma and territorial volume, cell density as well as temporal changes in visual acuity, contrast sensitivity and macroscopic brain morphology is based on the number of animals per group. In a very few cases, extreme outliers that were more than 3 standard deviations away from the respective group mean were excluded from further statistical analysis. Statistical analysis of MRI data is described in detail in section 2.5 (Deformation based morphometry). Data are reported as means with SEM or SD.

### 2.14 Data and Code availability

The data that support the findings of this study as well as the MATLAB code used for the morphometric analyses are available from the corresponding author upon reasonable request.

## 3. Results

### 3.1 Improved vision with open eye following MD

Brain plasticity induced by MD can be studied behaviorally by using optometry (Prusky et al., 2004): Animals watching moving vertical sine wave gratings track the grating with reflexive head movements. This tracking behavior depends on the ability of the animals to visually identify the grating, which in turn depends on the spatial frequency and the contrast of the grating. In non-deprived control animals, the VA of both eyes was stable during the entire observation period (p = 1, RM ANOVA on Ranks, n = 10, data not shown). In left-eye-deprived animals, VA of the contralateral OE increased in an exponential manner to reach a maximum of 37.0 ± 1.7 % above BL at day 9 post-MD (p < 0.001, RM ANOVA on Ranks, Fig. 1B). The rate of improvement of this visual perception learning, as represented by the daily increase in VA, was greatest at day one post-MD and then gradually decreased to zero (Fig. 1C). Similar increases in visual function were also observed for CS (ranging from +9.5 ± 0.3 % up to +14.4 ± 0.6 % at day 9 post-MD, depending on the spatial frequency of the grating, p < 0.001, RM ANOVA on Ranks, Fig. 1D+E, supplementary table S1).

The plasticity of VA was use-dependent: it increased to a larger extent in animals that underwent repetitive optometry (+37.0 ± 1.7 % at day 9 post-MD, p < 0.001, RM ANOVA on Ranks, n = 24, Fig. 1B) compared with animals without repetitive optometric testing (+24.6 ± 0.7 % at day 10 post-MD, p < 0.001, RM ANOVA on Ranks, n= 10, data not shown), thus indicating that daily testing by itself supported plasticity.

### 3.2 Macro-structural brain plasticity in visual and lateral entorhinal cortex

It has been shown that the MD-induced increases of VA and CS were associated with neuronal plasticity in V1B as well as in subcortical circuits (Prusky et al., 2006). Next, we evaluated MRI data by using DBM in a longitudinal study design to determine whether and where MD was accompanied by macro-structural brain plasticity. The DBM tool that we used allowed for detection of circumscribed brain volume changes that were not restricted to predefined regions of interest (Gaser et al., 2012; Herrmann et al., 2012) and the results reported here are based on identified clusters as shown in Fig. 2A. A series of four three-dimensional T2-weighted MRI image data sets was acquired for each animal, whereas the first data set was measured immediately before MD to obtain an individual BL image (Fig. 1A). The remaining three data sets were acquired sequentially at days 3, 7 and 10 post-MD. First, we screened the entire brain for dynamic volume changes reflecting the time course of daily visual improvement rates by testing for an initial tissue swelling. Compared with controls, MD resulted in a transient increase in grey matter volume at day 3 post-MD in the middle layers of the V1B contra-OE by 6.1 ± 1.1 %, and this was followed by a subsequent decline by 3.7 ± 1.1 % beyond the control levels at day 10 post-MD (RM ANOVA, threshold of p < 0.05, FWE corrected, Fig. 2A+B, supplementary table S2). The altered spatial information received by the animals with MD has to be processed in the hippocampus; hence, a swelling of the grey matter was also apparent in all layers of the LEnt contra-OE, which is the relay station into the hippocampus, by 6.4 ± 1.2 % (RM ANOVA, threshold of p < 0.05, FWE corrected, Fig. 2A+C, supplementary table S2). In line with visual improvement rates, this swelling regressed towards control levels at day 10 post-MD.

**Figure 2.**
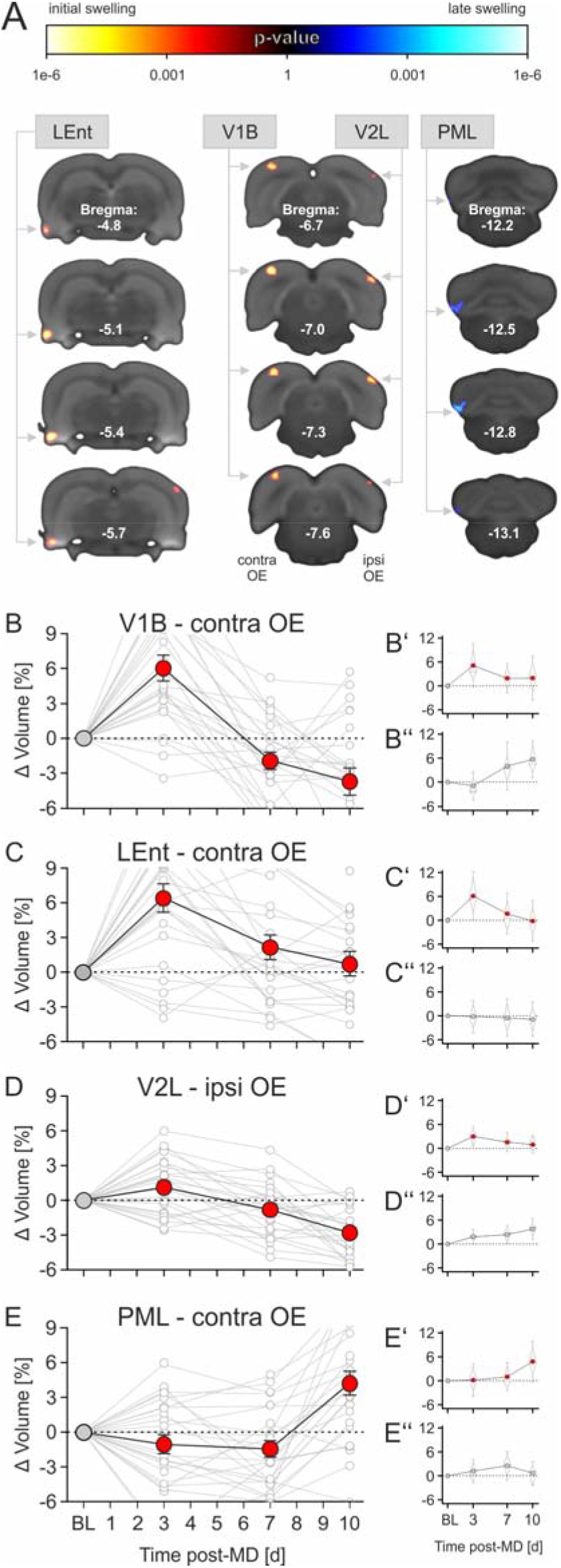
MD-induced macro-structural brain plasticity. **(A)** Statistical parametric maps demonstrating MD-induced temporal changes of brain morphology. Results of the RM ANOVA analyzing qualitative interactions between time and group (at an uncorrected threshold of p < 0.001) are superimposed on coronal sections of the reference brain. Yellow colors show regions where the brain in the MD-group (n = 24) initially swells in contrast to controls (n = 23). Vice versa, blue colors show brain regions where the volume in the MD-group shows a gain over time (late swelling). Significant clusters were found in the primary binocular visual cortex (V1B) and the lateral entorhinal cortex (LEnt) in the hemisphere contra-OE (threshold of p < 0.05, FWE corrected). The clusters in the paramedian lobule of the cerebellum (PML, contra-OE) and in the lateral secondary visual cortex (V2L, ipsi-OE) reached significance at an uncorrected threshold of p < 0.001. Numbers in slices indicate the position relative to Bregma. For detailed values and statistics, see supplementary table S2. **(B-E)** Quantification of MD-induced percentage changes in volume (Δ Volume) normalized to the mean of controls. Fast transient swelling of around 6 % in V1B and LEnt on day 3 post-MD is most prominent. Data are presented as mean ± sem. For better visualization, y-axes are truncated to the same values and some individual data points are hidden. The small plots in **B’-E’** and **B”-E”**, respectively, show the mean of the quantitative data separately for the MD group (red dots) and the controls (grey dots). Boxes indicate the 25th and 75th percentile, the line indicates the median, whiskers indicate sd.

The volume of the cerebellar paramedian lobulus (PML, contra-OE) increased with a delay at day 10 post-MD (RM ANOVA, uncorrected threshold of p < 0.001), thus possibly indicating motor adjustment to the altered visual experience. Small brain volume changes were also detected in the lateral part of the secondary visual cortex (V2L, ipsi-OE, RM ANOVA, uncorrected threshold of p < 0.001, Fig. 2A+D,E, supplementary table S2), possibly related to an increased multisensory signal processing (Hirokawa et al, 2008).

We further tested for brain volume changes on the basis of *a priori* selection of brain areas. Masks created with the brain atlas were superimposed on the brain images (Gaser et al., 2012) including V1B and LEnt. Comparing images made prior to and at day 3 post-MD confirmed volume changes in LEnt and V1B contra-OE (data not shown).

In the present study, the main effects on brain volume were observed in V1B and LEnt contra-OE. Several studies have shown that MD also causes plasticity in the visual cortex contralateral to the closed eye. These studies often perform an analysis after reopening of the eye, comparing electrical activity or intrinsic optical signals in the visual cortices after stimulation of the eye which remained open to signals after stimulation of the eye which underwent deprivation (cf. Hofer et al., 2006). Before MD, visually evoked neuronal activity in V1B is dominated by stimulation from the contralateral eye, whereas MD results in a shift towards a symmetric response rate (Rose et al., 2016; Sato and Stryker, 2008). With this paradigm, ocular dominance in the cortex contralateral to the eye which remained open is not significantly shifted by MD (Greifzu et al., 2011; Sato and Stryker, 2008). In our MRI-based analysis, neither the unbiased approach nor the *a priori* analysis of the MRI data revealed a change in brain volume in the primary visual cortex contralateral to the closed eye. When comparing these results one has to keep in mind that the paradigms employed are distinctly different.

### 3.3 No cell proliferation associated with macro-structural brain plasticity

Next, we used histology in a cross-sectional approach to analyse whether cell proliferation contributes to MD-induced brain volume changes (Fig. 3A). We focused on V1B and LEnt, and the AU was selected as control region. BrdU was used to label the nuclei of newborn cells in the first 3 days post-MD. Compared with controls, the density of BrdU+ cells did not increase; instead, there was a non-significant decrease in the density in the hemisphere contra-OE at day 3 post-MD in LEnt by 8.8 ± 4.7 % (p: 0.430, Student T-test) and in V1B by 7.8 ± 4.3 % (p: 0.196, Student T-test). This reduction was not observed in the hemisphere ipsi-OE (LEnt: p: 0.956, one outlier excluded from the MD-3d group; V1B: p: 0.894, Student T-test) and was also not present in the AU control region (contra-OE: p: 0.983, ipsi-OE: p: 0.930, Fig. 3C-E, for detailed data and statistics see supplementary table S3). To further confirm these results, we additionally determined the density of S100B+ astrocytes. The density of S100B+ cells decreased transiently in the hemisphere contra-OE at day 3 post-MD, by 7.9 ± 1.8 % in LEnt (F_(2,17)_: 6.45, p: 0.008, 1way ANOVA, Tukey HSD) and by 7.4 ± 1.2 % in V1B (F_(2,20)_: 5.83, p: 0.01, 1way ANOVA, Tukey HSD), and returned to normal at day 10 post-MD. No changes were observed in the corresponding brain areas ipsi-OE (LEnt: F_(2,18)_: 0.268, p: 0.768, one outlier excluded from the MD-10d group; V1B: F_(2,16)_: 0.007, p: 0.993, 1way ANOVA, Tukey HSD) and in AU (contra-OE: F_(2,19)_: 0.028, p: 0.972; ipsi-OE: F_(2,18)_: 0.271, p: 0.766, 1way ANOVA, Tukey HSD, Fig. 3F-H, for detailed data and statistics see supplementary table S3). We conclude that cell proliferation (of neurons or glial cells) does not contribute to grey matter swelling under the present paradigm. The reduction in the densities of BrdU+ and S100B+ cells was in the same range and was compatible with a “dilution” caused by the observed 6 −7 % increase in grey matter volume (cf. Vernon et al., 2014).

**Figure 3.**
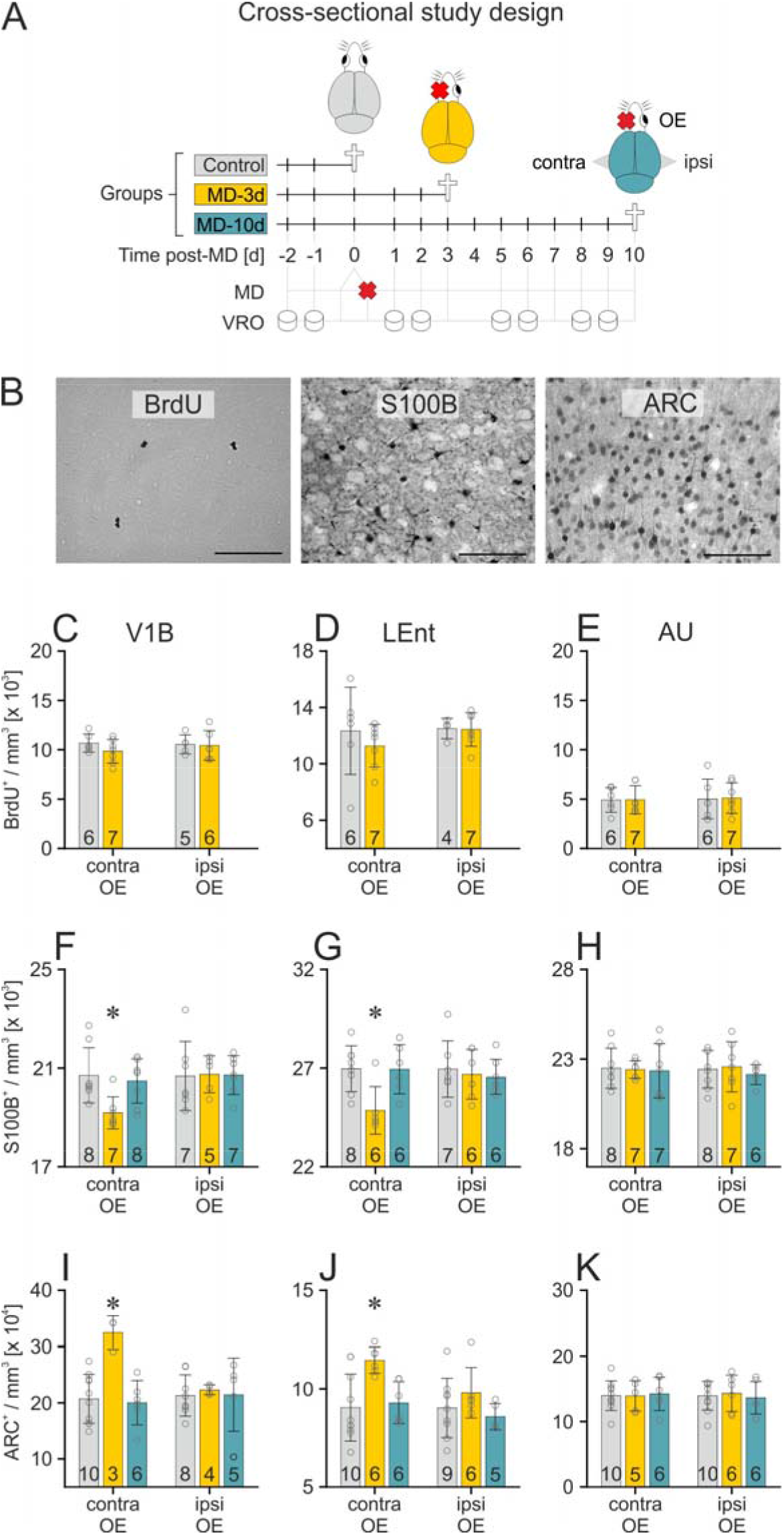
Stereological analyses in brain areas with macro-structural plasticity. **(A)** Cross-sectional study design: brains were analyzed without MD (control, grey) as well as at day 3 (yellow) or day 10 (blue) post-MD. **(B)** Representative brain sections showing BrdU+, S100B+ and ARC+ cells as used for stereological analyses, scale bar: 100 μm. **(C-E)** BrdU+ cells showed a non-significant decrease in thedensity in V1B and LEnt contra-OE at day 3 post-MD. **(F-H)** Density of S100B+ cells was transiently decreased at day 3 post-MD in V1B and LEnt contra-OE, with no alterations in AU. (I-K) Density of ARC+ cells was transiently increased at day 3 post-MD in V1B and LEnt contra-OE, with no alterations in AU. Data are presented as the mean ± sd. Numbers in bar-graphs show the amount of animals per group (n). *p < 0.05 vs. control, 1-way ANOVA, Tukey HSD. For detailed values and statistics, see supplementary table S3.

### 3.4 Synaptic plasticity associated with macro-structural brain plasticity

Expression of ARC is believed to be crucial for all known forms of synaptic plasticity (Korb and Finkbeiner, 2011). We therefore analyzed ARC expression using immunohistochemistry. Compared with controls, the density of ARC+ cells increased transiently over time in the hemisphere at day 3 post-MD in LEnt contra-OE by 26.6 ± 3.0 % (F _(2,19)_: 6.5, p: 0.007, 1way ANOVA, Tukey HSD) and by 56.8 ± 8.3 % in V1B contra-OE (F_(2,16)_: 10.9, p: 0.001, 1way ANOVA, Tukey HSD). It declined towards control levels at day 10 post-MD. The density of ARC+ cells remained unaltered in LEnt ipsi-OE (F _(2,17)_: 1.3, p: 0.303, 1way ANOVA, Tukey HSD) and V1B ipsi-OE (F_(2,14)_: 0.08, p: 0.925, one outlier excluded from the MD-10d group, 1way ANOVA, Tukey HSD) and bilaterally in AU (contra-OE: F_(2,18)_: 0.03, p: 0.970, one outlier excluded from the MD-3d group; ipsi-OE: F_(2,19)_: 0.12, p: 0.891, 1way ANOVA, Tukey HSD, Fig. 3I-K, for detailed data and statistics see supplementary table S3). Thus, the number of synaptically plastic neurons increased in the brain areas that display volume changes in MRI, whereas the brain areas ipsi-OE did not show such an increase.

### 3.5 Growth of neuronal dendrites in brain areas with macro-structural plasticity

A recent study has identified increased density of dendritic spines as a cellular correlate of changes in brain structure detected by post-mortem voxel-based morphometry (Keifer et al., 2015). We therefore utilized Golgi-stained neurons to analyze the structure of pyramidal dendrites and their spine density in V1B, LEnt and AU. We analyzed basal dendrites, since they allowed a definite attribution to pyramidal cells. Furthermore, basal dendrites receive input from layer 4 and also directly from the thalamus (Petreanu et al., 2009; Hooks et al, 2011) and have been reported to show a more pronounced use-dependent plasticity than apical dendrites (Seaton et al., 2020). Compared with controls, the lengths of 1^st^ order basal pyramidal dendrites remained unchanged in all brain areas examined (for detailed data and statistics see supplementary table S4), except for an small increase over time in AU ipsi-OE (F_(2,53)_: 3.835, p: 0.028, 1way ANOVA, Tukey HSD, Fig. 4B-D). In contrast, the lengths of the 2^nd^–3^rd^ order basal pyramidal dendrites increased over time at day 3 post-MD in LEnt contra-OE by 42.3 ± 8.5 % (F_(2,60)_: 8.94, p < 0,001, 1way ANOVA, Tukey HSD) and bilaterally in V1B by 56.8 ± 8.2 % contra-OE (F_(2,60)_: 19,165, p < 0.001, 1way ANOVA, Tukey HSD) and by 42.3 ± 6.9 % ipsi-OE (F_(2,49)_: 13.26, p < 0.001, 1way ANOVA, Tukey HSD), thus indicating a pronounced structural plasticity of 2^nd^–3^rd^ order basal dendrites of pyramidal neurons.

**Figure 4.**
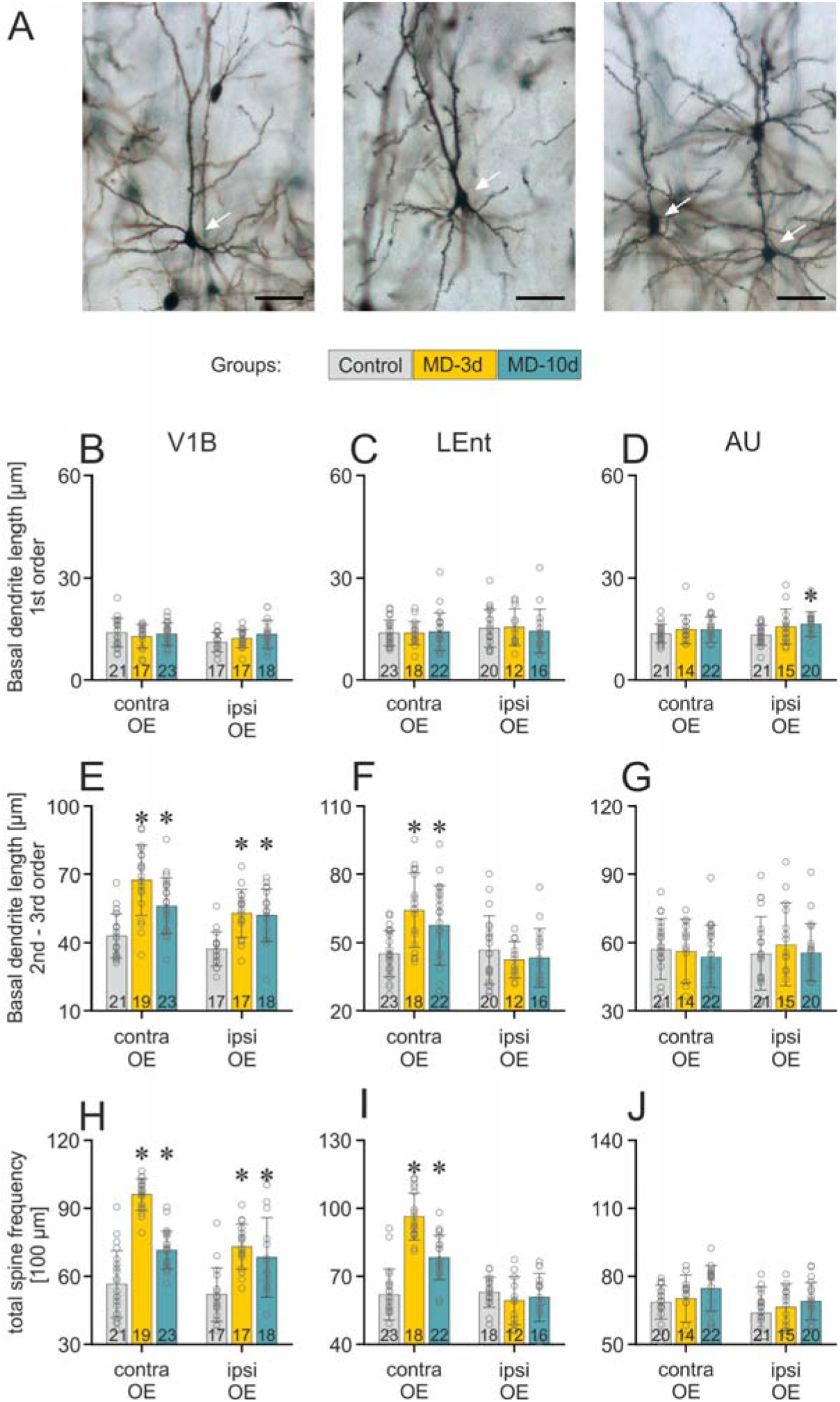
MD-induced neuronal structural changes in brain areas with macro-structural plasticity. **(A)** Representative Golgi-stained pyramidal neurons in V1B, LEnt and AU, scale bar: 50 μm. **(B-D)** Length of 1^st^ order basal dendrites was unaffected by MD, except for a small increase over time in AU ipsi-OE. **(E-G)** Length of 2^nd^–3^rd^ order basal dendrites was increased bilaterally in V1B and contra-OE in LEnt up to day 10 post-MD whereas no alterations were found in AU. **(H-J)** Total spine frequency on 2nd-3rd-order basal dendrites was increased bilaterally in V1B and contra-OE in LEnt up to day 10 post-MD whereas no alterations were found in AU. Data are presented as the mean ± sd. Numbers in bar-graphs show the amount of cells per group (N). *p < 0.05 vs. control, 1-way ANOVA, Tukey HSD. For detailed values and statistics, see supplementary tables S4 and S5A-C. Study design is shown in Fig. 3A, grey bars: control, yellow bars: day 3 post-MD, blue bars: day 10 post-MD.

In electron microscopic analyses, dendrites constitute about 20-30 % to grey matter volume, (Thomas et al., 2012). 1.6 % of the volume is contributed by 2^nd^–3^rd^ order basal dendrites of pyramidal neurons (Klenowski et al., 2017; Wang et al., 2018). A 1% increase in dendritic length results in a 0.75% increase in volume of basal dendrites (Gilman et al., 2017). Based on these estimations, the observed 42-57% increase in length of 2^nd^-3^rd^ order basal dendrites of pyramidal neurons in our study should produce a 0.50-0.67% increase in overall grey matter volume, accounting for 7.9-11% of macroscopic tissue swelling in LEnt and V1B observed with MRI.

Unlike the transient macroscopic grey matter volume increase observed via MRI, the dendritic length only partially regressed at day 10 post-MD: it remained elevated by 27.7 ± 8.2 % in LEnt contra-OE (F_(2,60)_: 8.94, p < 0,001, 1way ANOVA, Tukey HSD), by 30.6 ± 5.9 % in V1B contra-OE (F_(2,60)_: 19,165, p < 0.001, 1way ANOVA, Tukey HSD) and by 40.0 ± 7.3 % in V1B ipsi-OE (F_(2,49)_: 13.26, p < 0.001, 1way ANOVA, Tukey HSD). In AU, which served as a control region, the length of pyramidal 2^nd^–3^rd^ order dendrites did not change (contra-OE: F_(2,54)_: 0.321, p: 0.727; ipsi-OE: F_(2,53)_: 0.322, p: 0.726, 1way ANOVA, Tukey HSD); similarly, it remained unaltered in LEnt ipsi-OE (F_(2,45)_: 0.508, p: 0.605, 1way ANOVA, Tukey HSD, Fig. 4E-G, for detailed data and statistics see supplementary table S4).

### 3.6 Massive proliferation of dendritic spines in brain areas with macro-structural plasticity

Using the same Golgi stains, we next determined the spine frequency on 2^nd^–3^rd^ order dendrites (Fig. 4H-J, supplementary table S5A-C). Compared with controls, total spine frequency was increased over time at day 3 post-MD in LEnt contra-OE by 55.5 ± 3.9 % (F_(2,60)_:54.34, p < 0.001, 1way ANOVA, Tukey HSD), and bilaterally in V1B by 69.8 ± 2.8 % contra-OE (F_(2,60)_: 68.5, p < 0.001, 1way ANOVA, Tukey HSD) and by 40.8 ± 4.6 % ipsi-OE (F_(2,49)_: 12.76, p < 0.001, 1way ANOVA, Tukey HSD). Similar to the expansion of the 2^nd^–3^rd^ order dendrites, this increase partially persisted at day 10 after MD, with an elevation of 26.3 ± 3.4 % in LEnt contra-OE (F_(2,60)_: 54.34, p < 0.001, 1way ANOVA, Tukey HSD) and increases of 26.5 ± 3.1 % (F_(2,60)_: 68.5, p < 0.001, 1way ANOVA, Tukey HSD) and 31.6 ± 8.0 % (F_(2,49)_: 12.76, p < 0.001, 1way ANOVA, Tukey HSD) in V1B contra-OE and ipsi-OE, respectively. In AU, the number of spines was bilaterally stable over time (contra-OE: F_(2,53)_: 1.715, p: 0.19; ipsi-OE: F_(2,53)_: 1.347, p: 0.269, 1way ANOVA, Tukey HSD). Together, the increase in dendrite length in association with an increase in the number of spines per section of dendrite indicated an impressive structural plasticity of neurons.

Next, we classified five types of spines on the basis of their length (small: ≤ 1 μm, medium: 1 −1.5 μm and large: 1.5 - 4.5 μm) and shape (mushroom and spiny). The latter are typically characteristic of a mature state that is occupied with synapses (Kasai et al., 2010) (Fig. 5A). At day 3 post-MD, the numbers of “young” spines (small, medium, large) as well as of “mature” spines (mushroom and spiny) increased in the range of 34.0 ± 8.9 % to 83.9 ± 5.9 % per 100 μm of dendrite length. At day 10 post-MD, when the animals had reached a new stable VA, and grey matter volume had been reduced towards normal levels, the number of mature spines remained elevated by nearly the same extent as at day 3 post-MD in LEnt contra-OE. In contrast, the relative increase in the number of mature spines regressed towards BL in V1B contra-OE (Fig. 5B). No changes in spine frequency were found in AU (Fig.5B-D, supplementary table S5A-C).

**Figure 5.**
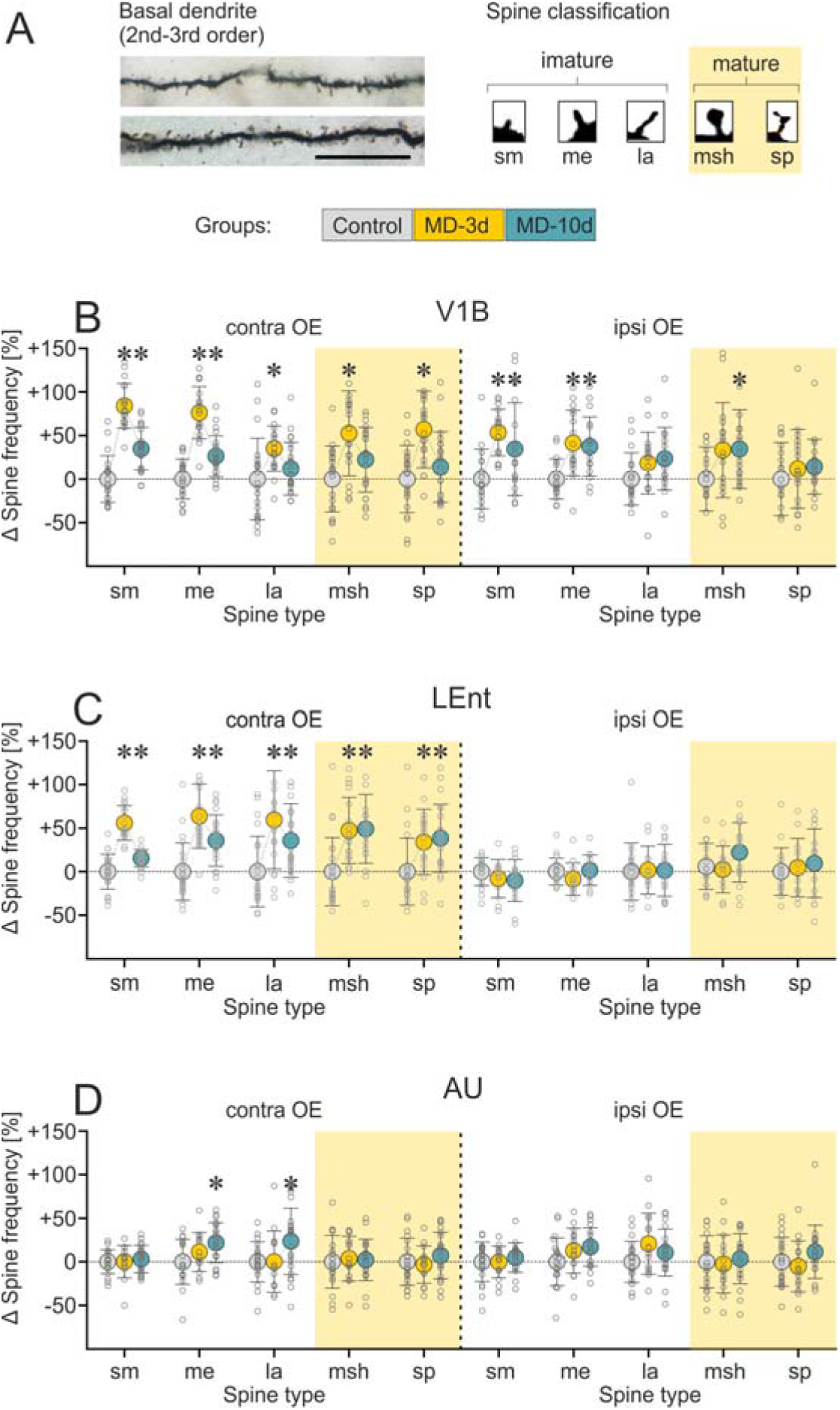
MD-induced spine plasticity in brain areas with macro-structural plasticity. **(A)** Representative images of Golgi-stained 2^nd^–3^rd^ order basal dendrites (scale-bar: 25 μm) and the spine classification scheme to separate immature from mature spines (sm: small, me: medium, la: large, msh: mushroom, sp: spiny). **(B-D)** MD-induced increase in the frequency of mature spines was mainly restricted to the hemisphere contra-OE in which it was transient in V1B and permanent in LEnt, with no alterations in AU. Data are presented as the mean ± sd. *p < 0.05 vs. control, 1-way ANOVA, Tukey HSD (based on number of cells, N=12-23). For detailed values and statistics, see supplementary table S5A-C. Study design is shown in Fig. 3A, grey dots: control, yellow dots: day 3 post-MD, blue dots: day 10 post-MD.

According to electron microscopic analyses, dendritic spines constitutes 5% volume of grey matter (Chklovskii et al., 2002). Spines on 2^nd^-3^rd^ order basal dendrites of pyramidal neurons specifically contribute 0.35% to the grey matter volume (Klenowski et al., 2017; Wang et al., 2018). A 1% increase in spine density causes a 1.17 % increase in spine volume (Keifer et al., 2015). Based on these estimations, the 55.5-69.8% increase in spine density of 2^nd^-3^rd^ order basal dendrites observed in the present study would cause a 0.22-0.28 % increase in cortical volume and contribute 3.5-4.6 % to the cortical expansion in LEnt and V1B observed with MRI.

### 3.7 Reorganization of astrocyte complexity and size in brain areas with macro-structural plasticity

The length of neuronal dendrites and the number of neuronal spines remained partially elevated at day 10 after MD, while the macroscopic grey matter swelling as well as ARC expression regressed back towards BL. Synaptic activation, which induces long-term potentiation, causes a transient increase in the motility of perisynaptic astrocytic domains and is predictive of spine plasticity (Bernardinelli et al., 2014), and astrocytic processes may be necessary for the establishment of stable new synapses (Allen and Lyons, 2018; Farhy-Tselnicker et al., 2017; Kim et al., 2018; Singh et al., 2016; Sudhof, 2018). Therefore, we analysed astrocyte complexity and volume in the macro-structurally plastic brain areas after MD.

The number of GFAP-stained primary astrocytic processes was unaffected by MD in LEnt (contra-OE: F_(2,123)_: 1.1, p: 0.343; ipsi-OE: F_(2,122)_: 0.46, p: 0.633, 1way ANOVA, Tukey HSD), V1B (contra-OE: F_(2,123)_: 0.567, p: 0.569; ipsi-OE: F_(2,123)_: 0,083, p: 0.92, 1way ANOVA, Tukey HSD) and AU (contra-OE: F_(2,123)_: 0.2, p: 0.824; ipsi-OE: F_(2,123)_: 0.11, p: 0.899,1way ANOVA, Tukey HSD, Fig. 6B-D, supplementary table S6). Compared with controls, the complexity of processes was decreased 3 days post-MD by 15.1 ± 0.8 % in LEnt contra-OE (F_(2,123)_: 35.4, p < 0.001, 1way ANOVA, Tukey HSD), and by 13.2 ± 4.6 % (F_(2,123)_: 31, p < 0,001, 1way ANOVA, Tukey HSD) and 9.6 ± 3.3 % (F_(2,123)_: 20.86, p < 0,001, 1way ANOVA, Tukey HSD) in V1B contra-OE and ipsi-OE, respectively. At day 10 post-MD, this decrease changed into an increase of 13.4 ± 4.1 % in LEnt contra-OE (F_(2,123)_: 35.4, p <0.001, 1way ANOVA, Tukey HSD) and of 21.8 ± 4.6 % (F_(2,123)_: 31, p < 0,001, 1way ANOVA, Tukey HSD) and 16.9 ± 3.4 % (F_(2,123)_: 20.86, p < 0,001, 1way ANOVA, Tukey HSD) in V1B contra-OE and ipsi-OE, respectively. No changes in GFAP-stained astrocytic processes were observed in AU (contra-OE: F_(2,123)_: 0.26, p: 0.770; ipsi-OE: F_(2,123)_: 0.4, p: 0.396, 1way ANOVA, Tukey HSD) and LEnt ipsi-OE (F_(2,123)_: 1.8, p: 0.171, 1way ANOVA, Tukey HSD, Fig. 6E-G, supplementary table S6). GFAP is a cytoskeletal protein that helps to maintain the mechanical shape and strength of astrocytes. It is present in large and intermediate astrocytic processes but not in small processes (Reichenbach et al., 2010). We hypothesized that a breakdown in GFAP polymerization allows a structural reorganization of astrocytic processes with increased motility, which is then consolidated with increased complexity by a new polymerization of GFAP filaments.

**Figure 6.**
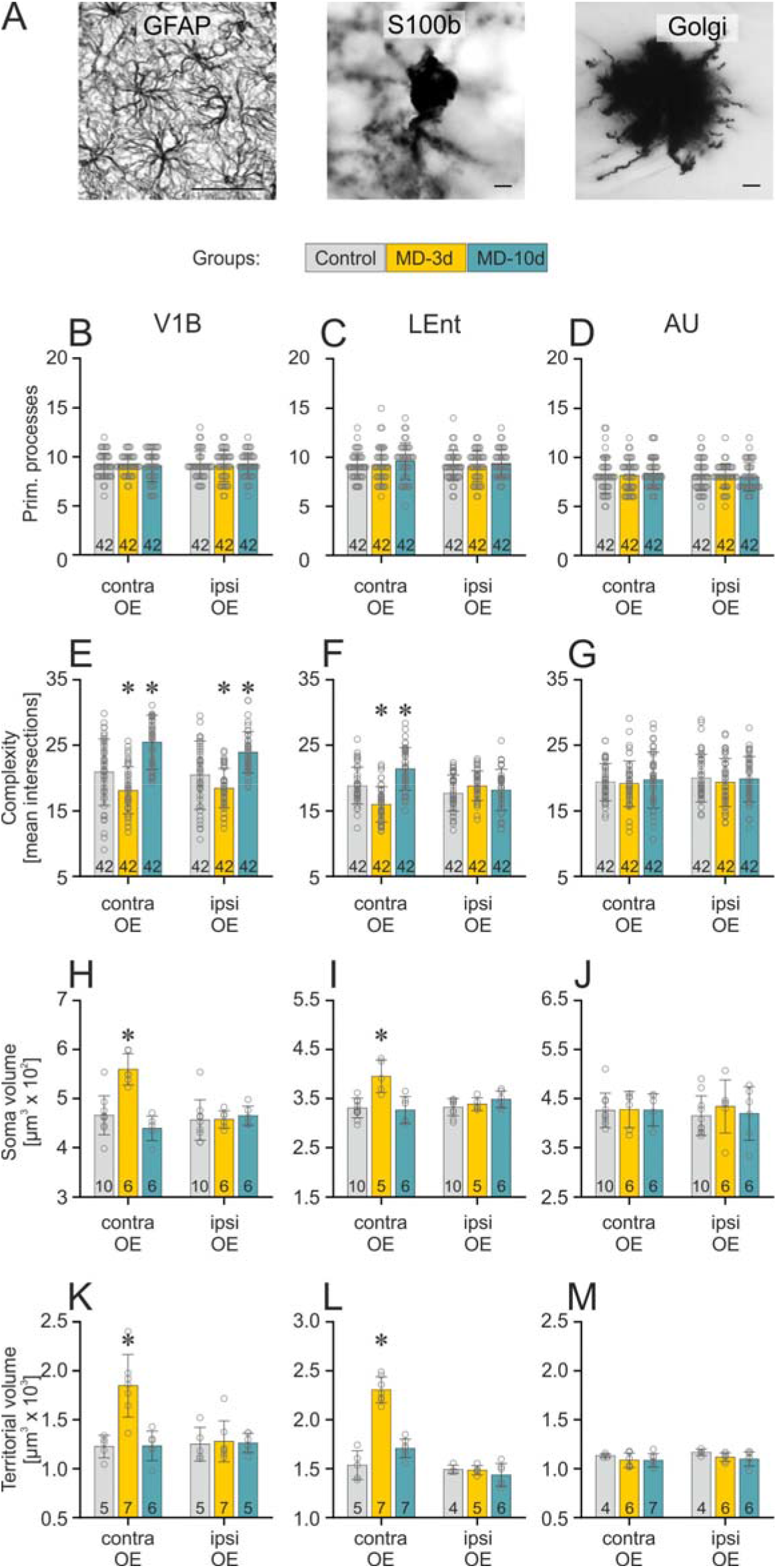
Reorganization of astrocytes in brain areas with macro-structural plasticity. **(A)** Representative GFAP+, S100+ and Golgi-stained astrocytes used to estimate cytoskeleton complexity, soma- and territorial volume, scale bars: 50 μm (GFAP) and 10μm (S100, Golgi). **(B-D)** Numbers of primary astrocytic processes were unaffected by MD in LEnt, V1B and AU. **(E-G)** Complexity of the astrocytic cytoskeleton was changed bilaterally in V1B and contra-OE in LEnt with a decrease at day 3 followed by an increase at day 10 post-MD. No alterations were found in AU. **(H-J)** Astrocytic soma volume was transiently increased at day 3 post-MD contra-OE in V1B and LEnt. No alterations were found in AU. **(K-M)** Astrocytic territorial volume was transiently increased at day 3 post-MD contra-OE in V1B and LEnt. No alterations were found in AU. Data are presented as the mean ± sd, using statistics for astrocytic complexity based on number of cells (N, shown as numbers in bar-graphs) and for astrocytic soma and territorial volume based on number of animals (n, shown as numbers in bar-graphs). *p < 0.05, 1-way ANOVA, Tukey HSD. For detailed values and statistics, see supplementary tables S6 and S7. Study design is shown in Fig. 3A, grey bars: control, yellow bars: day 3 post-MD, blue bars: day 10 post-MD.

To obtain a more complete measure of the territorial volume of astrocytes (the volume of neuropil occupied by a single astrocyte), we used a modified Golgi staining protocol that specifically impregnates astrocytes but not neurons (Grosche et al., 2013; Gull et al., 2015). Compared with controls, astrocytic territorial volumes increased at day 3 post-MD by 50.2 ± 3.3 % in LEnt (F_(2,16)_: 67.1, p < 0.001, 1way ANOVA, Tukey HSD) and by 50.7 ± 9.9 % in V1B contra-OE (F_(2,15)_: 15.69, p < 0.001, 1way ANOVA, Tukey HSD) and recovered at day 10 post-MD. No changes in territorial volumes were observed in V1B (F_(2,15)_: 0.044, p: 0.957, 1way ANOVA, Tukey HSD) and LEnt (F_(2,12)_: 0.7, p: 0.516, 1way ANOVA, Tukey HSD) in the hemisphere ipsi-OE or bilaterally in AU (contra-OE: F_(2,14)_: 0.82, p: 0.461; ipsi-OE: F_(2,13)_: 1.8, p: 0.194, 1way ANOVA, Tukey HSD, Fig. 6K-L, supplementary table S7). Similarly, astrocyte soma volume was assessed on S100B+ cells and revealed a transient increase at day 3 post-MD in the hemisphere contra-OE both in LEnt (+19.5 ± 4.5 %, F_(2,18)_: 15.8, p < 0.001, 1way ANOVA, Tukey HSD) and in V1B (+20.1 ± 2.8 %, F_(2,19)_: 21, p < 0.001, 1way ANOVA, Tukey HSD), which returned to control levels at day 10 post-MD. No changes in astrocytic soma volume were observed in V1B (F_(2,19)_: 0,156, p: 0.856, 1way ANOVA, Tukey HSD) and LEnt (F_(2,18)_: 1.8, p: 0.195, 1way ANOVA, Tukey HSD) in the hemisphere ipsi-OE or in AU bilaterally (contra-OE: F_(2,19)_: 0.005, p: 0.995; ipsi-OE: F_(2,19)_: 0.3, p: 0.750, 1way ANOVA, Tukey HSD, Fig. 6H-J, supplementary table S7).

We then calculated the extent to which the observed changes in astrocyte volume contributed to the macroscopic grey matter swelling in LEnt and V1B at day 3 post-MD. Through electron microscopic analyses, it has been estimated that the fraction of astrocytes in cortical grey matter volume is approximately 9 % (Genoud et al., 2006; Jones and Greenough, 1996; Thomas et al., 2012). On the basis of this value, a 50 % increase in astrocytic volume explains 71-74 % of the macroscopic tissue swelling in LEnt and V1B, thus suggesting that a major proportion of MD-induced grey matter swelling observed via MRI in the present experiments was associated by astrocyte plasticity.

## 4. Discussion

When the constant stream of information from one eye is lost, the brain adapts by processing more information from the preserved eye. In the present paradigm, this results in an increased visual acuity of the non-deprived eye within the first three days. Volumetric MRI revealed strong adaptations in V1B contra-OE; unlike in humans this receives more than 95% of corresponding retinal ganglion cell projections in rats (Dreher et al., 1985). Adaptations also appeared in several other brain areas, especially the entorhinal cortex contra-OE, which conveys visual information to the hippocampus (Nau et al., 2018). Part of the observed structural plasticity – i.e. the late adaptation in the cerebellum – may be an indirect consequence of an adaptive motor behavior.

The pattern of experience-dependent structural brain plasticity observed in the present study differed between brain areas. In LEnt contra-OE a pronounced increase in brain volume detected by MRI was associated with synaptic plasticity as indicated by ARC staining, maturation of dendritic spines and a strong transient increase in the densities of spines. The increase in brain volume vanished within 10 days, however, the spines partially matured and persisted 10 days after onset of monocular deprivation of the ipsilateral eye. A pattern similar to that in LEnt was observed in the V1B contra-OE, although this area lacked the persistent increase in the number of mature dendritic spines.

The neuronal plasticity observed with the present protocol was large: the length of 2^nd^–3^rd^ order basal pyramidal dendrites increased by 40 – 60 %, and spine density by 50 to 70 % in LEnt and V1B contra-OE at day 3 post-MD. A similar positive correlation between MRI-detectable volume changes with markers of neuronal process remodeling could also be shown by Lerch et al. (2011). In our case, in LEnt contra-OE more than 50% of the new spines persisted and matured after 10 days, thus resulting in a longer term neuronal structural plasticity. This brain area thus showed a *gain of structure plasticity*.

After onset of monocular deprivation, the visual acuity increased continuously up to about day 5 (repetitive use-dependent plasticity over several days). In this period there were many new morphologically young spines on the neurons in the brain areas showing grey matter swelling. To which extent these new, morphologically young spines constitute a receptive state for the establishment of new synapses has to be addressed in further studies (Constantinidis and Klingberg, 2016; Xu et al., 2009).

In contrast to LEnt and V1B contra-OE, the visually deprived V1B ipsi-OE displayed no macro-structural plasticity of grey matter detectable in the MRI, no change in grey matter volume on the microscopic level, no transcriptional activity as indicated by ARC staining, no persistent increase in the number of mature spines, and no increase in the soma or territorial volume of astrocytes. As a sign of structural plasticity, an increase in neuronal dendrite length and young spines was observed and was associated with a restructuring of astrocyte processes, as indicated by GFAP expression. The plasticity in this brain area is usually attributed to “*homeostatic plasticity*” (Keck et al., 2017).

Cell proliferation did not contribute to the DBM signal. Rather, a reduced cell density was measured during grey matter swelling, compatible with an extension of the territory between the nuclei of neurons, as well as astrocytes. In AU which served as control region in the present study no change in cell proliferation, no structural and functional plasticity in neurons, no restructuring of astrocytic processes or somata, and no changes of territorial volume of astrocytes were observed.

Our data showed synaptic plasticity and spine proliferation on 2^nd^-3^rd^ order basal dendrites of pyramidal neurons with grey matter volume expansion in LEnt and V1B contra-OE at day 3 post-MD. In the present study first order dendrites remained unchanged. Based on data from literature we calculated that the observed structural plasticity of 2^nd^-3^rd^ order basal dendrites and their spines increases the grey matter volume by about 1%. This may escape detection in our experimental arrangement. We should, however, emphasize that we did not analyze the contribution of dendritic elements distal to 2^nd^-3^rd^ order dendrites, and that this are rather indirect estimations.

According to the present data, the grey matter volume changes measured by MRI were dominated by structural changes in astrocytes. A transient increase in astrocytic territorial volume was seen in LEnt and V1B contra-OE; the time course of these volume changes correlated with time course of macro-structural grey matter plasticity as well as with transient volume changes derived from the distribution of S100 beta stained astrocytic somata. Astrocytic territorial volume increased by 50 %, with an initial reduction and subsequent increase in GFAP expression in intermediate and ramified processes indicating a restructuring of these processes. Degree and time course of astrocyte reorganization fit well to the reversible increase in brain volume as measured by MRI or astrocyte density.

Recent studies have emphasized the roles of perisynaptic astrocytic processes in the establishment and the stabilization of new synapses (Araque et al., 1999; Pannasch et al., 2014; Santello et al., 2019). Astrocytes span local territories (Bushong et al., 2002) and communicate with one another via mechanisms such as calcium waves (Cornell-Bell et al., 1990; Poskanzer and Yuste, 2016). Large-scale astrocyte reorganization, which involves both distal and proximal processes and even the cell body, may thus generate a permissive environment for synaptic plasticity in the affected astrocytic territories (Adamsky et al., 2018). Sagi et al. (2012) found after short-term water maze navigation changes in diffusion metrics associated with changes in astrocytes morphometry. Woo et al. (2018) recently reported volumetric brain plasticity in the hippocampus of humans and mice which required aquaporin-4 channels in astrocytes; aquaporin-4 knockdown impaired volumetric plasticity and long term potentiation induced by theta burst. Notably, human astrocytes, compared with rodent astrocytes, are more complex and appear to better support synapse reorganization (Oberheim et al., 2006).

In the present study, macroscopic plasticity of brain structure as determined by MRI was associated with neuronal dendrite and spine plasticity and astrocyte reorganization. The macroscopic brain plasticity observed in our study was transient, leaving the question open whether pronounced repetitive learning may finally result in persistently enhanced volumes of certain brain areas, as suggested in studies on taxi drivers, or musicians (Gaser and Schlaug, 2003; Maguire et al., 2000).

## 5. Conclusion

We conclude that learning may be associated with pronounced structural rearrangements of brain grey matter. *Astrocyte plasticity* may contribute substantially to this use-dependent brain plasticity, which has long been regarded as a domain governed nearly exclusively by neurons.

## Acknowledgments

The authors received support from grants from BMBF (Bernstein Focus, 01GQ0923 to O.W.W), BMBF (JenAge, 0315581 to O.W.W), BMBF (Irestra, 16SV7209 to O.W.W), DFG (HHDP, FOR 1738, WI 830/10-2, 830/12-1 to O.W.W), TMWWDG (ProExellenz, RegenerAging-FSU-I-03/14 to O.W.W).

## Author contributions

O.W.W., S.S. and S.G. designed the experiments with contributions from all authors, O.W.W. supervised the study, S.S. and S.G. performed experiments, M.B., A.I., A.U. participated in behavioral and molecular measurements, K.H.H. and J.R.R. established and optimized MRI protocols and participated in the experiments, S.S. and S.G. analysed functional and molecular data, C.G. established volumetric mouse analysis tool, C.K., C.G. and S.S. analysed volumetric data, O.W.W., S.S. and S.G. interpreted the data and wrote the paper with contributions from all authors.

## Supplemental Information

**Supplementary table S1:**
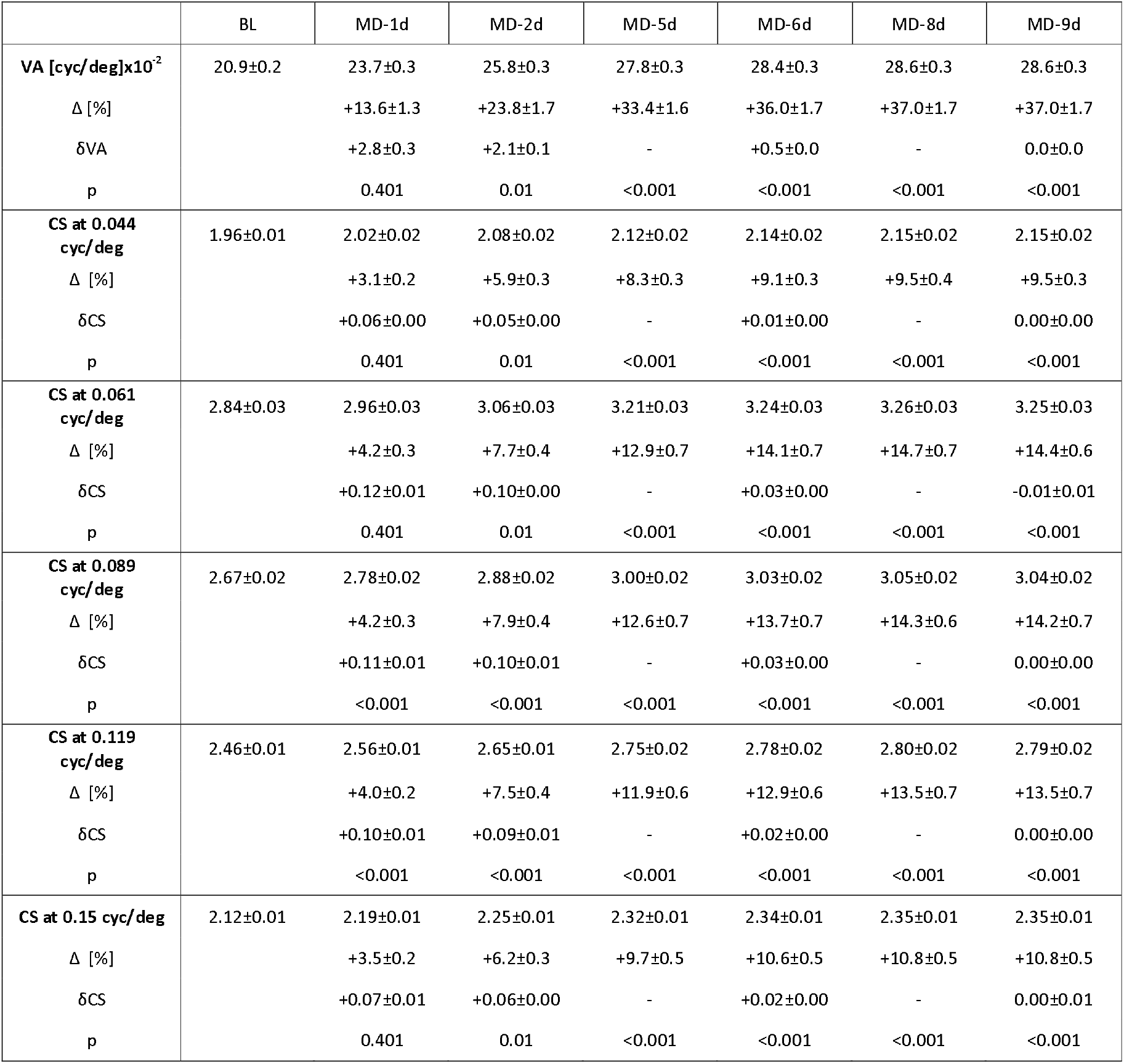
Visual acuity (VA) and contrast sensitivity (CS) examined by optometry represented as mean ± sem; BL: baseline, MD-1d to −10d: time-points of optometry following MD, Δ shows the percental difference vs. BL, δ shows the daily improvement rates; statistics: RM ANOVA on Ranks (n=24). Study design is shown in Fig. 1A.

**Supplementary table S2:** DBM results. Areas showing MD-induced volume changes. RM ANOVA at a threshold of p < 0.05, FWE corrected and at a threshold of p < 0.001 (uncorrected), t-test for volume changes between MD-3d and MD-10d in MD animals (n = 24) compared to controls (n = 23). LEnt: lateral entorhinal cortex, V1B: primary visual cortex, binocular area, V2L: secondary visual cortex, lateral part, PML: paramedian lobule of the cerebellum. Study design is shown in Fig. 1A.

| Area | Hemisphere | Coordinates |  |  | p <0.05(FWE) | p <0.001(uncorrected) |  |
| --- | --- | --- | --- | --- | --- | --- | --- |
|  |  | X | Y | Z | p value | p value | cluster size |
| LEnt | contra-OE | -6.7 | -8.4 | -5.4 | 0.011 | 0.006 | 82 |
| V1B | contra-OE | -4.5 | -2.0 | -7.0 | 0.011 | 0.004 | 95 |
| V2L | ipsi-OE | 6.1 | -2.9 | -7.2 | - | 0.011 | 70 |
| PML | contra-OE | -5.1 | -6.4 | -12.6 | - | 0.032 | 46 |

**Supplementary table S3:**
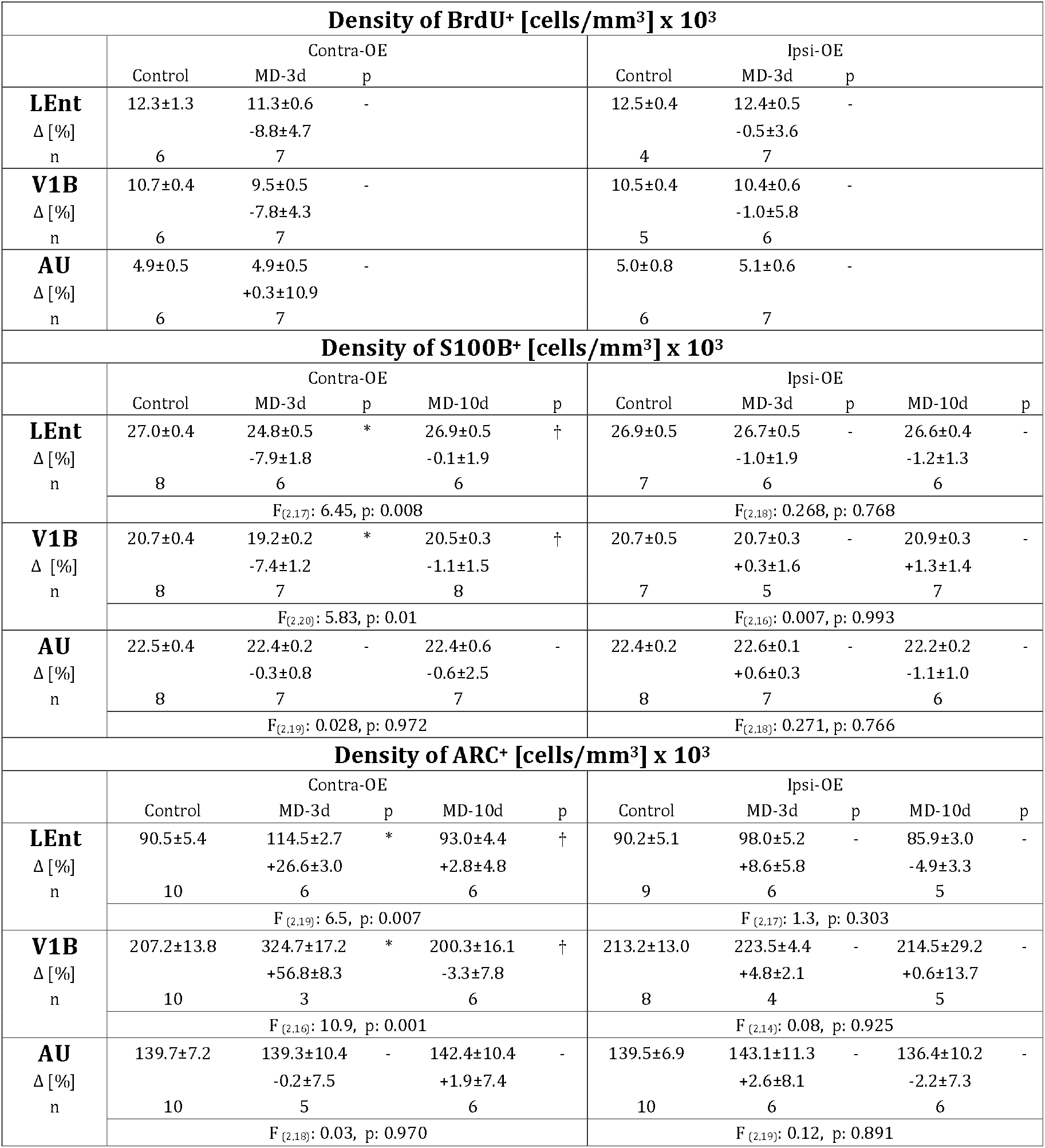
Density of BrdU+, S100B+ and ARC+ cells represented as mean ± sem; LEnt: lateral entorhinal cortex, V1B: primary visual cortex, binocular area, AU: primary auditory cortex; Δ shows the percental difference vs. controls, n = number of animals; * p MD vs. control < 0.05, † p MD-10d vs. MD-3d < 0.05; for BrdU+ student-t test and for S100B+, ARC+ 1way ANOVA, Tukey HSD was used. Study design is shown in Fig. 3A.

**Supplementary table S4:** Basal pyramidal dendrite length presented as mean ± sem, using statistics based on number of cells; LEnt: lateral entorhinal cortex, V1B: primary visual cortex, binocular area, AU: primary auditory cortex, Δ shows the percental difference vs. controls; N=number of cells, n=number of animals; * p MD vs. Control < 0.05; † p MD-10d vs. MD-3d < 0.05; 1way ANOVA, Tukey HSD. Study design is shown in Fig. 3A.

| Basal dendrite length [μm] | | | | | | | | | | |
| --- | --- | --- | --- | --- | --- | --- | --- | --- | --- | --- |
|  | Contra-OE |  |  |  |  | Ipsi-OE |  |  |  |  |
|  | Control | MD-3d | p | MD-10d | p | Control | MD-3d | p | MD-10d | p |
| **LEnt** | 14.0±0.8 | 13.9±0.8 | - | 14.3±1.2 | - | 15.3±1.2 | 17.7±1.6 | - | 14.5±1.6 | - |
| Δlength [%] | | -0.6±5.7 | | +2.2±8.5 | | | +2.2±10.1 | | -5.2±10.5 | |
| N/n | 23/7 | 18/5 |  | 22/6 |  | 20/7 | 12/3 |  | 16/5 |  |
| 1. order | F<sub>(2,60)</sub>: 0.036, p: 0.964 |  |  |  |  | F<sub>(2,45)</sub>: 0.172, p: 0.843 |  |  |  |  |
| **LEnt** | 45.3±2.1 | 64.4±3.5 | * | 57.8±3.7 | * | 46.9±3.4 | 42.6±2.3 | - | 43.5±3.2 | - |
| Δlength [%] | | +42.3±8.5 | | +27.7±8.2 | | | -9.1±5.0 | | -7.3±6.9 | |
| N/n | 23/7 | 18/5 |  | 22/6 |  | 20/7 | 12/3 |  | 16/5 |  |
| 2.-3. order | F<sub>(2,60)</sub>: 8.94, p < 0.001 |  |  |  |  | F<sub>(2,45)</sub>: 0.508, p: 0.605 |  |  |  |  |
| **V1B** | 13.9±0.9 | 12.8±0.8 | - | 13.5±0.7 | - | 11.1±0.7 | 12.1±0.6 | - | 13.4±1.0 | - |
| Δlength [%] | | -8.0±6.0 | | -2.9±5.0 | | | +9.8±5.9 | | +21.0±8.7 | |
| N/n | 21/6 | 17/6 |  | 23/7 |  | 17/6 | 17/6 |  | 18/6 |  |
| 1. order | F<sub>(2,60)</sub>: 0.0708, p: 0.925 |  |  |  |  | F<sub>(2,49)</sub>: 2.198, p: 0.122 |  |  |  |  |
| **V1B** | 43.0±2.1 | 67.4±3.5 | * | 56.1±2.5 | *† | 37.2±1.8 | 52.9±2.9 | * | 52.1±2.7 | * |
| Δlength [%] | | +56.8±8.2 | | +30.6±5.9 | | | +42.3±6.9 | | +40.0±7.3 | |
| N/n | 21/6 | 19/6 |  | 23/7 |  | 17/6 | 17/6 |  | 18/6 |  |
| 2.-3. order | F<sub>(2,60)</sub>: 19.165, p<0.001 |  |  |  |  | F<sub>(2,49)</sub>: 13.26, p<0.001 |  |  |  |  |
| **AU** | 13.7±0.6 | 15.0±1.1 | - | 14.9±0.8 | - | 13.3±0.6 | 15.8±1.3 | - | 16.6±0.8 | * |
| Δlength [%] | | +9.1±8.2 | | +8.6±5.9 | | | +18.7±10.0 | | +24.2±6.1 | |
| N/n | 21/7 | 14/5 |  | 22/6 |  | 21/7 | 15/5 |  | 20/6 |  |
| 1. order | F<sub>(2,54)</sub>: 0.771, p: 0.467 |  |  |  |  | F<sub>(2,53)</sub>: 3.835, p: 0.028 |  |  |  |  |
| **AU** | 57.2±2.9 | 56.3±3.8 | - | 53.9±2.9 | - | 55.2±3.5 | 59.2±4.7 | - | 55.7±2.8 | - |
| Δlength [%] | | -1.5±6.6 | | -5.7±5.1 | | | +7.2±8.6 | | +0.8±5.1 | |
| N/n | 21/7 | 14/5 |  | 22/6 |  | 21/7 | 15/5 |  | 20/6 |  |
| 2.-3. order | F<sub>(2,54)</sub>: 0.321, p: 0.727 |  |  |  |  | F<sub>(2,53)</sub>: 0.322, p: 0.726 |  |  |  |  |

**Supplementary table S5A:**
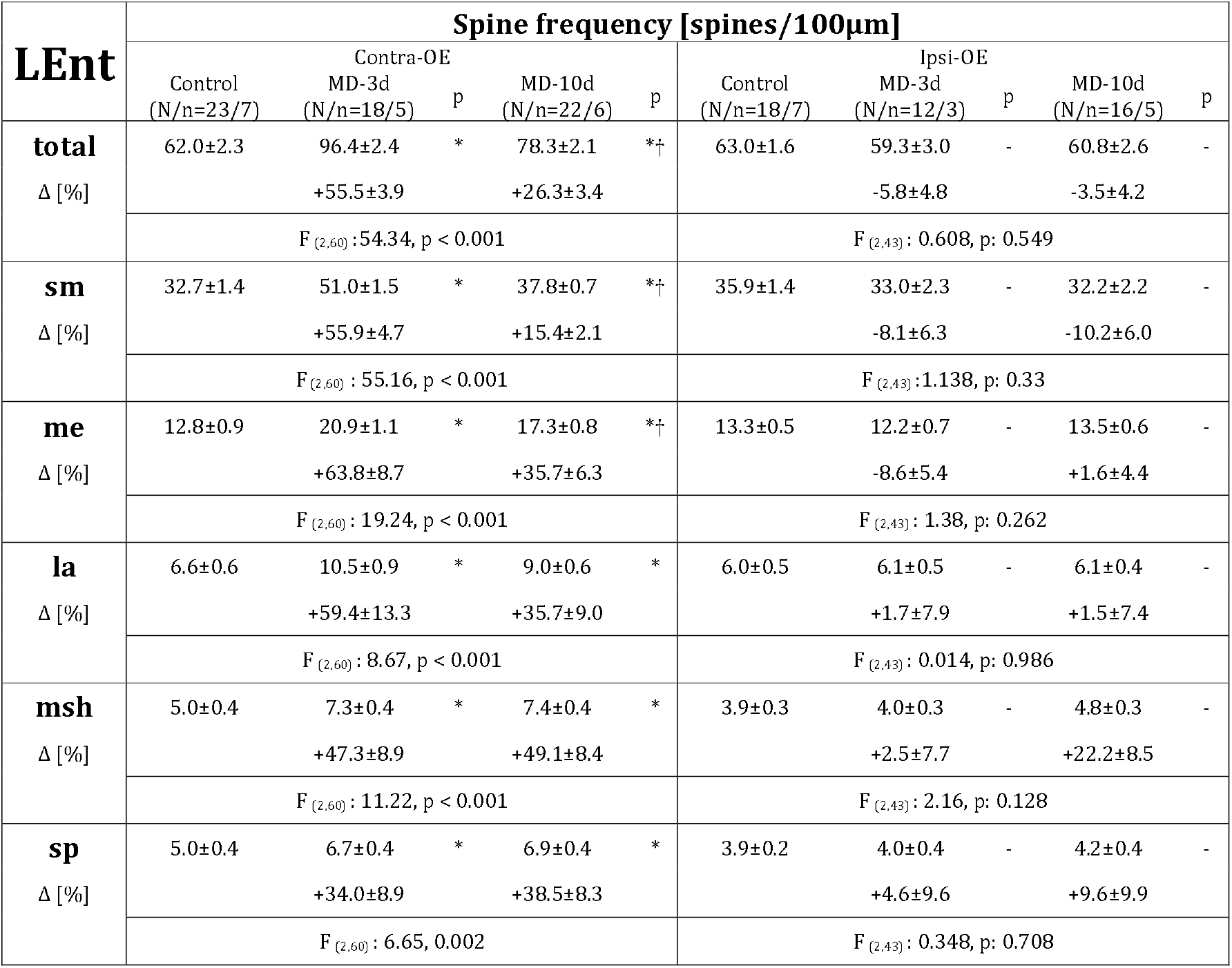
Spine frequency (spines per 100μm) on 2^nd^–3^rd^ order basal pyramidal dendrites in LEnt (lateral entorhinal cortex) presented as mean ± sem, using statistics based on number of cells; total: whole spine pool, sm: small spines, me: medium spines, la: large spines, msh: mushroom spines, sp: spiny spines, Δ shows the percental difference vs. controls; N=number of cells, n=number of animals; * p MD vs. control < 0.05; † p MD-10d vs. MD-3d < 0.05; 1way ANOVA, Tukey HSD. Study design is shown in Fig. 3A.

| LEnt | Spine frequency [spines/100µm] |  |  |  |  |  |  |  |  |  |
| --- | --- | --- | --- | --- | --- | --- | --- | --- | --- | --- |
|  | Contra-OE |  |  |  |  | Ipsi-OE |  |  |  |  |
|  | Control<br>(N/n=23/7) | MD-3d<br>(N/n=18/5) | p | MD-10d<br>(N/n=22/6) | p | Control<br>(N/n=18/7) | MD-3d<br>(N/n=12/3) | p | MD-10d<br>(N/n=16/5) | p |
| **total** | 62.0±2.3 | 96.4±2.4 | * | 78.3±2.1 | *† | 63.0±1.6 | 59.3±3.0 | - | 60.8±2.6 | - |
| Δ [%] |  | +55.5±3.9 |  | +26.3±3.4 |  |  | -5.8±4.8 |  | -3.5±4.2 |  |
|  | F<sub>(2,60)</sub>: 54.34, p < 0.001 |  |  |  |  | F<sub>(2,43)</sub>: 0.608, p: 0.549 |  |  |  |  |
| **sm** | 32.7±1.4 | 51.0±1.5 | * | 37.8±0.7 | *† | 35.9±1.4 | 33.0±2.3 | - | 32.2±2.2 | - |
| Δ [%] |  | +55.9±4.7 |  | +15.4±2.1 |  |  | -8.1±6.3 |  | -10.2±6.0 |  |
|  | F<sub>(2,60)</sub>: 55.16, p < 0.001 |  |  |  |  | F<sub>(2,43)</sub>: 1.138, p: 0.33 |  |  |  |  |
| **me** | 12.8±0.9 | 20.9±1.1 | * | 17.3±0.8 | *† | 13.3±0.5 | 12.2±0.7 | - | 13.5±0.6 | - |
| Δ [%] |  | +63.8±8.7 |  | +35.7±6.3 |  |  | -8.6±5.4 |  | +1.6±4.4 |  |
|  | F<sub>(2,60)</sub>: 19.24, p < 0.001 |  |  |  |  | F<sub>(2,43)</sub>: 1.38, p: 0.262 |  |  |  |  |
| **la** | 6.6±0.6 | 10.5±0.9 | * | 9.0±0.6 | * | 6.0±0.5 | 6.1±0.5 | - | 6.1±0.4 | - |
| Δ [%] |  | +59.4±13.3 |  | +35.7±9.0 |  |  | +1.7±7.9 |  | +1.5±7.4 |  |
|  | F<sub>(2,60)</sub>: 8.67, p < 0.001 |  |  |  |  | F<sub>(2,43)</sub>: 0.014, p: 0.986 |  |  |  |  |
| **msh** | 5.0±0.4 | 7.3±0.4 | * | 7.4±0.4 | * | 3.9±0.3 | 4.0±0.3 | - | 4.8±0.3 | - |
| Δ [%] |  | +47.3±8.9 |  | +49.1±8.4 |  |  | +2.5±7.7 |  | +22.2±8.5 |  |
|  | F<sub>(2,60)</sub>: 11.22, p < 0.001 |  |  |  |  | F<sub>(2,43)</sub>: 2.16, p: 0.128 |  |  |  |  |
| **sp** | 5.0±0.4 | 6.7±0.4 | * | 6.9±0.4 | * | 3.9±0.2 | 4.0±0.4 | - | 4.2±0.4 | - |
| Δ [%] |  | +34.0±8.9 |  | +38.5±8.3 |  |  | +4.6±9.6 |  | +9.6±9.9 |  |
|  | F<sub>(2,60)</sub>: 6.65, 0.002 |  |  |  |  | F<sub>(2,43)</sub>: 0.348, p: 0.708 |  |  |  |  |

**Supplementary table S5B:** Spine frequency (spines per 100μm) on 2^nd^–3^rd^ order basal pyramidal dendrites in V1B (primary visual cortex, binocular area) presented as mean ± sem, using statistics based on number of cells; total: whole spine pool, sm: small spines, me: medium spines, la: large spines, msh: mushroom spines, sp: spiny spines, Δ shows the percental difference vs. controls; N=number of cells, n=number of animals; * p MD vs. control < 0.05; † p MD-10d vs. MD-3d < 0.05; 1way ANOVA, Tukey HSD. Study design is shown in Fig. 3A.

| V1B | Spine frequency [spines/100µm] |  |  |  |  |  |  |  |  |  |
| --- | --- | --- | --- | --- | --- | --- | --- | --- | --- | --- |
|  | Contra-OE |  |  |  |  | Ipsi-OE |  |  |  |  |
|  | Control<br>(N/n=21/6) | MD-3d<br>(N/n=19/6) | p | MD-10d<br>(N/n=23/7) | p | Control<br>(N/n=17/6) | MD-3d<br>(N/n=17/6) | p | MD-10d<br>(N/n=18/6) | p |
| **total** | 56.6±3.2 | 96.0±1.6 | * | 71.5±1.7 | *† | 51.9±2.9 | 73.1±2.4 | * | 68.4±4.1 | * |
| Δ [%] |  | +69.8±2.8 |  | +26.5±3.1 |  |  | +40.8±4.6 |  | +31.6±8.0 |  |
|  | F<sub>(2,60)</sub>: 68.5, p < 0.001 |  |  |  |  | F<sub>(2,49)</sub>: 12.76, p < 0.001 |  |  |  |  |
| **sm** | 25.7±1.5 | 47.3±1.5 | * | 34.7±1.3 | *† | 25.1±2.1 | 38.5±1.6 | * | 33.7±3.1 | * |
| Δ [%] |  | +83.9±5.9 |  | +34.6±5.1 |  |  | +53.4±6.5 |  | +34.3±12.5 |  |
|  | F<sub>(2,60)</sub>: 57.72, p < 0.001 |  |  |  |  | F<sub>(2,49)</sub>: 9.08, p < 0.001 |  |  |  |  |
| **me** | 12.3±0.6 | 21.7±0.8 | * | 15.6±0.6 | *† | 11.7±0.7 | 16.5±1.1 | * | 16.0±0.9 | * |
| Δ [%] |  | +76.1±6.8 |  | +26.1±4.9 |  |  | +41.3±9.1 |  | +37.3±8.0 |  |
|  | F<sub>(2,60)</sub>: 52.63, p < 0.001 |  |  |  |  | F<sub>(2,49)</sub>: 8.76, p < 0.001 |  |  |  |  |
| **la** | 8.2±0.8 | 11.0±0.5 | * | 9.2±0.5 | † | 6.9±0.5 | 8.1±0.6 | - | 8.5±0.6 | - |
| Δ [%] |  | +35.0±5.9 |  | +11.9±6.3 |  |  | +18.2±8.6 |  | +23.3±8.5 |  |
|  | F<sub>(2,60)</sub>: 7.04, p: 0.002 |  |  |  |  | F<sub>(2,49)</sub>: 2.96, p: 0.061 |  |  |  |  |
| **msh** | 5.3±0.4 | 8.1±0.6 | * | 6.5±0.4 | † | 4.0±0.4 | 5.3±0.5 | - | 5.3±0.4 | * |
| Δ [%] |  | +52.5±11.2 |  | +22.2±7.7 |  |  | +33.2±13.2 |  | +34.4±10.6 |  |
|  | F<sub>(2,60)</sub>: 9.6, p < 0.001 |  |  |  |  | F<sub>(2,49)</sub>: 4.28, p: 0.019 |  |  |  |  |
| **sp** | 5.0±0.4 | 7.8±0.5 | * | 5.7±0.4 | † | 4.3±0.4 | 4.8±0.5 | - | 4.9±0.3 | - |
| Δ [%] |  | +56.8±10.1 |  | +13.8±8.4 |  |  | +11.6±11.0 |  | +14.0±7.4 |  |
|  | F<sub>(2,60)</sub>: 15.94, p < 0.001 |  |  |  |  | F<sub>(2,49)</sub>: 0.775, p: 0.466 |  |  |  |  |

**Supplementary table S5C:**
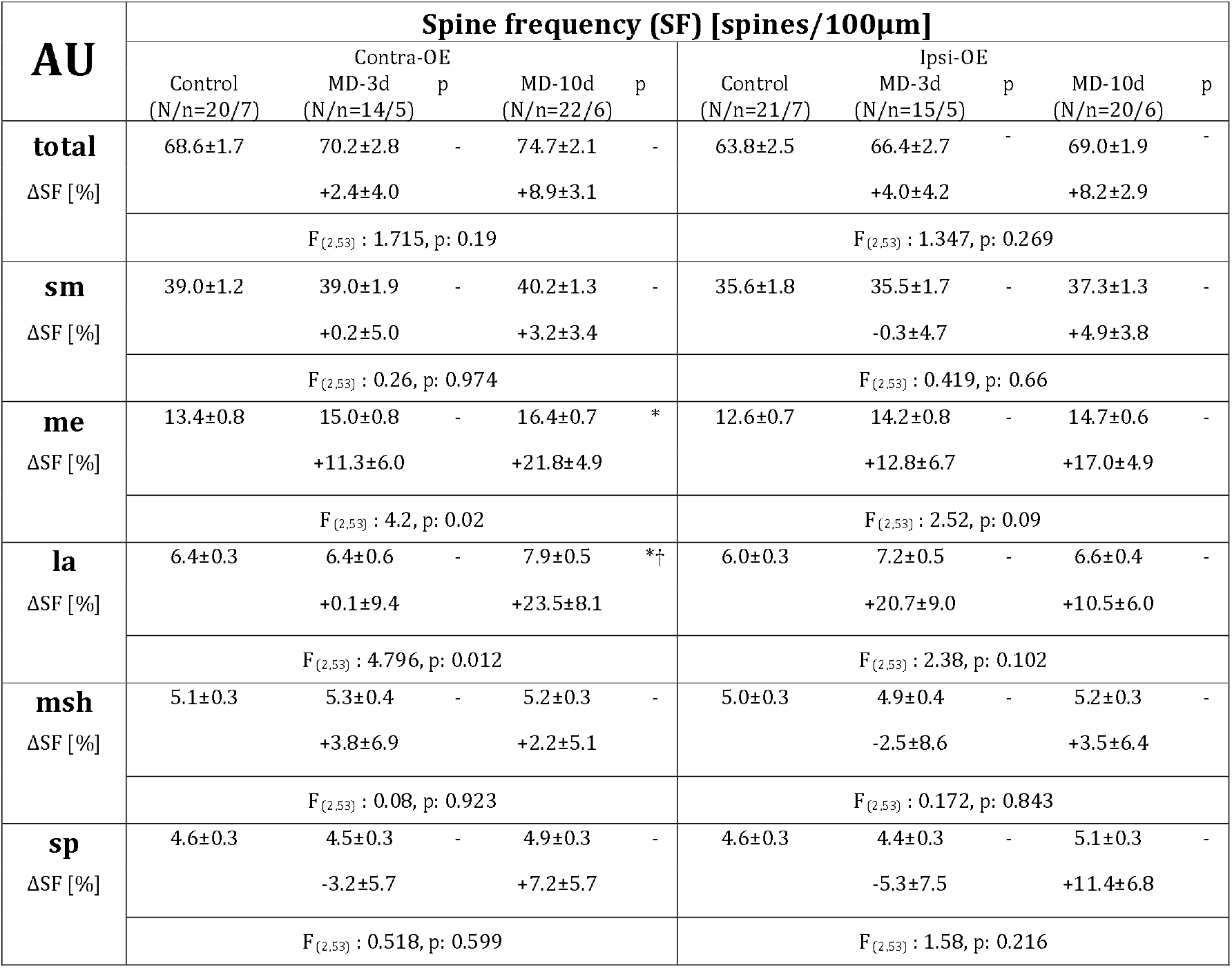
Spine frequency (SF, spines per 100μm) on 2^nd^–3^rd^ order basal pyramidal dendrites in AU (primary auditory cortex) presented as mean ± sem, using statistics based on number of cells; total: whole spine pool, sm: small spines, me: medium spines, la: large spines, msh: mushroom spines, sp: spiny spines, Δ shows the percental difference vs. controls; N=number of cells, n=number of animals; * p MD vs. control < 0.05; † p MD-10d vs. MD-3d < 0.05; 1way ANOVA, Tukey HSD. Study design is shown in Fig. 3A.

**Supplementary table S6:**
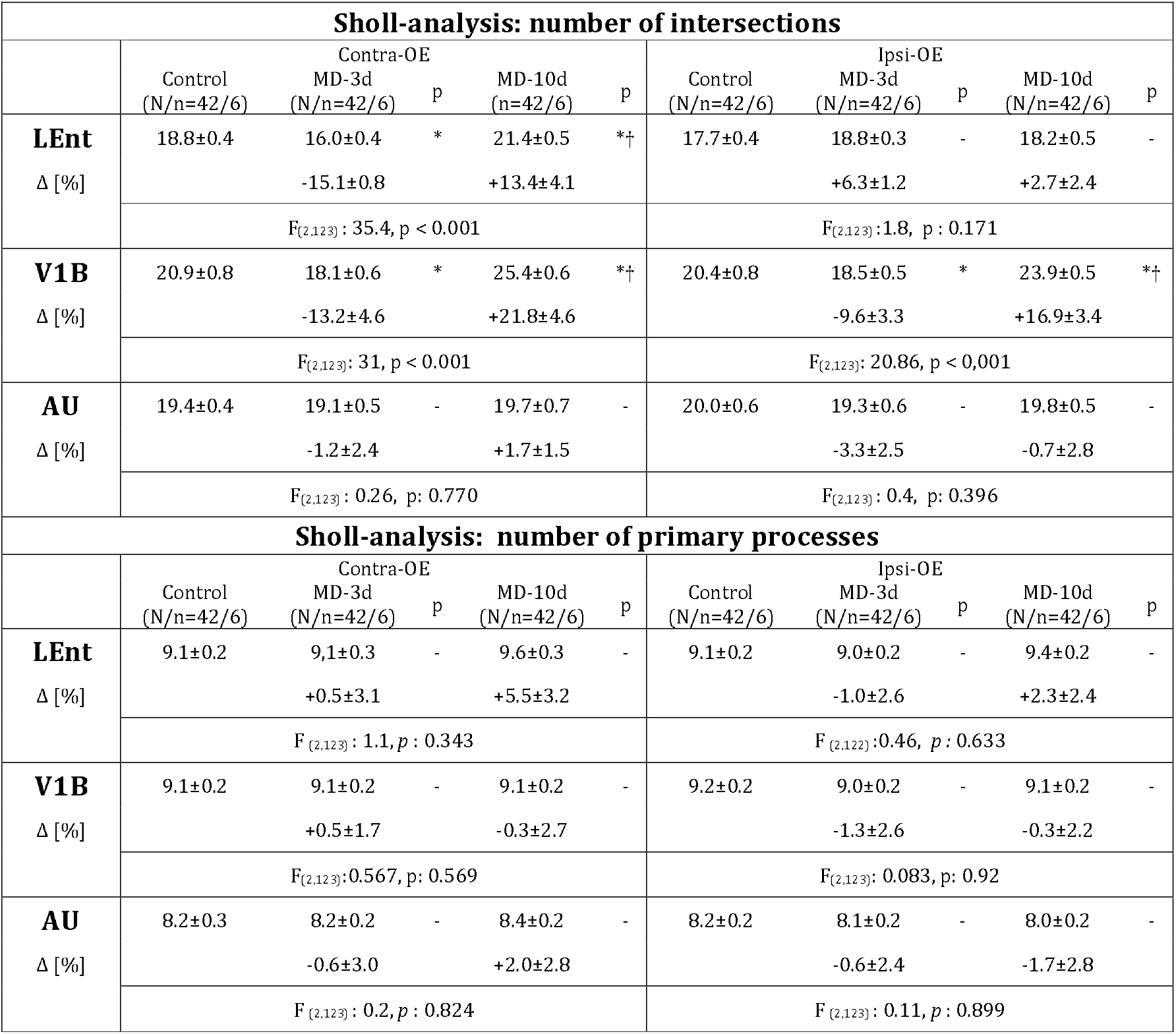
Astrocytic cytoskeleton complexity as analyzed by the Sholl’s concentric circle method. Number of intersections and number of primary processes are presented as mean ± sem, using statistics based on number of cells; LEnt: lateral entorhinal cortex, V1B: primary visual cortex, binocular area, AU: primary auditory cortex, Δ shows the percental difference vs. controls; N=number of cells, n=number of animals; * p MD vs. control < 0.05; † p MD-10d vs. MD-3d < 0.05; 1way ANOVA, Tukey HSD. Study design is shown in Fig. 3A.

**Supplementary table S7:** Astrocytic territorial volume and astrocytic soma volume are presented as mean ± sem; LEnt: lateral entorhinal cortex, V1B: primary visual cortex, binocular area, AU: primary auditory cortex, Δ shows the percental difference vs. controls; n=number of animals; * p MD vs. control < 0.05; † p MD-10d vs. MD-3d < 0.05; 1way ANOVA, Tukey HSD. Study design is shown in Fig. 3A.

| Astrocytic territorial volume in $\mu\text{m}^3 \times 10^2$ | | | | | | | | | | |
| --- | --- | --- | --- | --- | --- | --- | --- | --- | --- | --- |
|  | Contra-OE |  |  |  |  | Ipsi-OE |  |  |  |  |
|  | Control | MD-3d | p | MD-10d | p | Control | MD-3d | p | MD-10d | p |
| **LEnt** | 15.3±0.7 | 23.1±0.5 | * | 17.1±0.4 | † | 14.9±0.2 | 14.8±0.2 | - | 14.4±0.5 | - |
| $\Delta$ [%] | | +50.2±3.3 | | +11.2±2.3 | | | -0.6±1.4 | | -3.8±3.2 | |
| n | 5 | 7 |  | 7 |  | 4 | 5 |  | 6 |  |
|  | F<sub>(2,16)</sub>: 67.1, p < 0.001 |  |  |  |  | F<sub>(2,12)</sub>: 0.7, p: 0.516 |  |  |  |  |
| **V1B** | 12.3±0.5 | 18.5±1.2 | * | 12.3±0.6 | † | 12.5±0.8 | 12.8±0.8 | - | 12.6±0.4 | - |
| $\Delta$ [%] | | +50.7±9.9 | | +0.5±5.0 | | | +2.3±6.3 | | +1.1±3.2 | |
| n | 5 | 7 |  | 6 |  | 5 | 7 |  | 5 |  |
|  | F<sub>(2,15)</sub>: 15.69, p < 0.001 |  |  |  |  | F<sub>(2,15)</sub>: 0.044, p: 0.957 |  |  |  |  |
| **AU** | 11.3±0.1 | 10.9±0.3 | - | 10.8±0.3 | - | 11.7±0.2 | 11.2±0.2 | - | 11.0±0.3 | - |
| $\Delta$ [%] | | -4.0±2.7 | | -4.2±2.3 | | | -4.2±1.6 | | -5.9±2.6 | |
| n | 4 | 6 |  | 7 |  | 4 | 6 |  | 6 |  |
|  | F<sub>(2,14)</sub>: 0.82, p: 0.461 |  |  |  |  | F<sub>(2,13)</sub>: 1.8, p: 0.194 |  |  |  |  |
| Astrocytic soma volume in $\mu\text{m}^3 \times 10$ | | | | | | | | | | |
|  | Contra-OE |  |  |  |  | Ipsi-OE |  |  |  |  |
|  | Control | MD-3d | p | MD-10d | p | Control | MD-3d | p | MD-10d | p |
| **LEnt** | 33.1±0.6 | 39.5±1.5 | * | 32.7±1.1 | † | 33.2±0.6 | 33.9±0.6 | - | 34.8±0.7 | - |
| $\Delta$ [%] | | +19.5±4.5 | | -1.3±3.4 | | | +1.9±1.7 | | +4.8±2.1 | |
| n | 10 | 5 |  | 6 |  | 10 | 5 |  | 6 |  |
|  | F<sub>(2,18)</sub>: 15.8, p < 0.001 |  |  |  |  | F<sub>(2,18)</sub>: 1.8, p: 0.195 |  |  |  |  |
| **V1B** | 46.6±1.3 | 56.0±1.3 | * | 43.9±1.0 | † | 45.6±1.3 | 45.8±0.7 | - | 46.5±0.8 | - |
| $\Delta$ [%] | | +20.1±2.8 | | -5.7±2.2 | | | +0.3±1.6 | | +1.9±1.7 | |
| n | 10 | 6 |  | 6 |  | 10 | 6 |  | 6 |  |
|  | F<sub>(2,19)</sub>: 21, p < 0.001 |  |  |  |  | F<sub>(2,19)</sub>: 0.156, p: 0.856 |  |  |  |  |
| **AU** | 42.6±1.1 | 42.8±1.5 | - | 42.7±1.3 | - | 41.5±1.3 | 43.4±2.2 | - | 42.0±2.2 | - |
| $\Delta$ [%] | | +0.4±3.6 | | +0.2±3.1 | | | +4.5±5.3 | | +1.1±5.4 | |
| n | 10 | 6 |  | 6 |  | 10 | 6 |  | 6 |  |
|  | F<sub>(2,19)</sub>: 0.005, p: 0.995 |  |  |  |  | F<sub>(2,19)</sub>: 0.3, p: 0.750 |  |  |  |  |

